# Opposing regulatory logics converge on ABA receptors to govern the trade-off between growth and drought acclimation

**DOI:** 10.64898/2026.02.26.708209

**Authors:** Shuaikun Gu, Xiaoxiao Liu, Yuzi Zhang, Lei Zhang, Qingting Liu, Junyao Lu, Jinguang Huang, Guodong Yang, Kang Yan, Chengchao Zheng, Shizhong Zhang, Changai Wu

**Author notes:** **Corresponding authors:** Changai Wu; Shizhong Zhang. **E-mails:** Shuaikun Gu, Xiaoxiao Liu, Yuzi Zhang, Lei Zhang, Qingting Liu, Junyao Lu, Jinguang Huang, Guodong Yang, Kang Yan, Chengchao Zheng.

## Abstract

Plants dynamically balance growth with stress responses through abscisic acid (ABA). How ABA signaling, but how ABA sensitivity is finely tuned is unclear. Here, we uncover a regulatory network of leucine-rich repeat receptor-like kinases (LRR-RLKs) that directly controls ABA receptors stability. We identify GASSHO1 (GSO1) as a core component that phosphorylates the PYL2 and PYL4 receptors, targeting them for degradation and thus serving as a brake on ABA signaling. Under drought stress, ABA accumulates and suppresses expression of GSO1-activating CIF peptides, releasing this brake to increase PYLs and sensitivity. This mechanism opposes our previously identified CEPR2 pathway, where drought-induced CEP peptides inhibit the kinase to stabilize PYLs. The integration of these two antagonistic modules within an LRR-RLK network enables precise, dynamic control of ABA perception. This work reveals a molecular framework explaining how plants calibrate the critical balance between growth and drought acclimation.

## Introduction

The ability to perceive and respond to water deficit is fundamental to plant survival^1^. The phytohormone abscisic acid (ABA) serves as the primary chemical messenger coordinating these acclimatory responses^2–4^. The core ABA signaling pathway involves the binding of ABA to the receptors PYRABACTIN RESISTANCE 1 (PYR1) and its related proteins PYR1-like (PYL, also reported as REGULATORY COMPONENT OF ABA RECEPTORs [RCARs]), which subsequently inhibit clade A protein phosphatase 2Cs (PP2Cs), leading to the activation of SNF1-RELATED PROTEIN KINASE 2s (SnRK2s) and the phosphorylation of downstream effectors^5–10^. Although the abundance of PYL receptors is a critical determinant of signaling output, the upstream mechanisms that dynamically control PYL protein stability remain poorly understood.

Protein phosphorylation and targeted degradation represent a ubiquitous regulatory paradigm^11^. Our previous work identified one such pathway, whereby a leucine-rich repeat receptor-like kinase (LRR-RLK), C-TERMINALLY ENCODED PEPTIDE RECEPTOR 2 (CEPR2), phosphorylates PYL4, leading to its targeting for degradation in *Arabidopsis*. This phosphorylation of PYL4 is inhibited by the accumulation of C-TERMINALLY ENCODED PEPTIDEs (CEPs) encoded by genes whose expression is induced by ABA under drought stress^12,13^. The CEP and CEPR2 forms a typical negative feedback loop for PYL4.

The LRR-RLK family encompasses the largest plant-specific clade of the eukaryotic kinase superfamily, and over 200 members across various plant species^14^. Most LRR-RLKs localize to the plasma membrane and play roles for the the perception and transduction of diverse cellular signals. They control many plant developmental processes, such as CLAVATA 1 (CLV1) in meristem maintenance^15^, CLV3 INSENSITIVE RECEPTOR KINASE (CIK4) in stem cell fate^16^, BRASSINOSTEROID INSENSITIVE 1 (BRI1) and BRI1-ASSOCIATED RECEPTOR KINASE (BAK1) in brassinosteroid perception^17^, FLAGELLIN-SENSITIVE 2 (FLS2) in bacterial flagellin sensing^17^, IMPAIRED OOMYCETE SUSCEPTIBILITY 1 (IOS1) in ABA signal transduction^19,20^, GUARD CELL HYDROGEN PEROXIDE-RESISTANT 1 (GHR1) in stomatal closure^21^, and POLLEN RECEPTOR LIKE KINASE (PRK) in pollen tube development^22–24^. Given the considerable number of LRR-RLK family members, coupled with their established roles in sensing a broad spectrum of extracellular signals, we hypothesized that the regulatory framework governing PYL stability is more extensive than currently recognized. We therefore postulated that other LRR-RLK may potentially employ a distinct regulatory logic and converge onto PYL turnover, integrating a broad range of environmental and developmental cues into the ABA signaling network.

Among the LRR-RLKs in *Arabidopsis*, GASSHO1 (GSO1) is a compelling candidate for such a regulator. Initially characterized for its role in embryonic cuticle formation and root development, GSO1 is activated by the sulfated peptides CASPARIAN STRIP INTEGRITY FACTORs (CIFs), which promote the formation of a receptor complex and kinase activity^25–30^. However, a direct role for GSO1 in the core ABA signaling pathway remains undefined. Here, we conducted a systematic, large-scale screen to map the interaction landscape between 196 LRR-RLKs and the 14 ABA receptors of *Arabidopsis*, establishing an extensive regulatory network. We also performed a deep mechanistic dissection of one central node in this network, the GSO1–CIF module, uncovering a regulatory principle opposite to the established paradigm and revealing a sophisticated, multi-layered system for fine-tuning ABA perception.

## Results

### The interactome between LRR-RLKs and ABA receptors defines a crosstalk hub for signal integration

To systematically identify upstream regulators that could influence ABA receptor abundance, we performed a large-scale membrane-based yeast two-hybrid (MbY2H) screen, assaying 196 LRR-RLKs against PYR1 and the 13 PYL receptors (Supplementary Fig. 1). This screen identified 53 LRR-RLKs as potential interactors of at least one ABA receptor. Based on these results, we constructed a Membrane-based Interaction Network Database (MIND) (Fig. 1a). Analysis of the MIND revealed that PYR1 and nine PYLs, including PYL2, PYL4, and PYL5, serve as hubs, each associating with more than 25 of LRR-RLKs. These interacting LRR-RLKs are implicated in diverse signaling pathways related to development, environmental responses, and reproduction. The convergence of these pathways suggests that multiple LRR-RLK–PYL modules serve as central nodes for signal integration.

**Figure 1.**
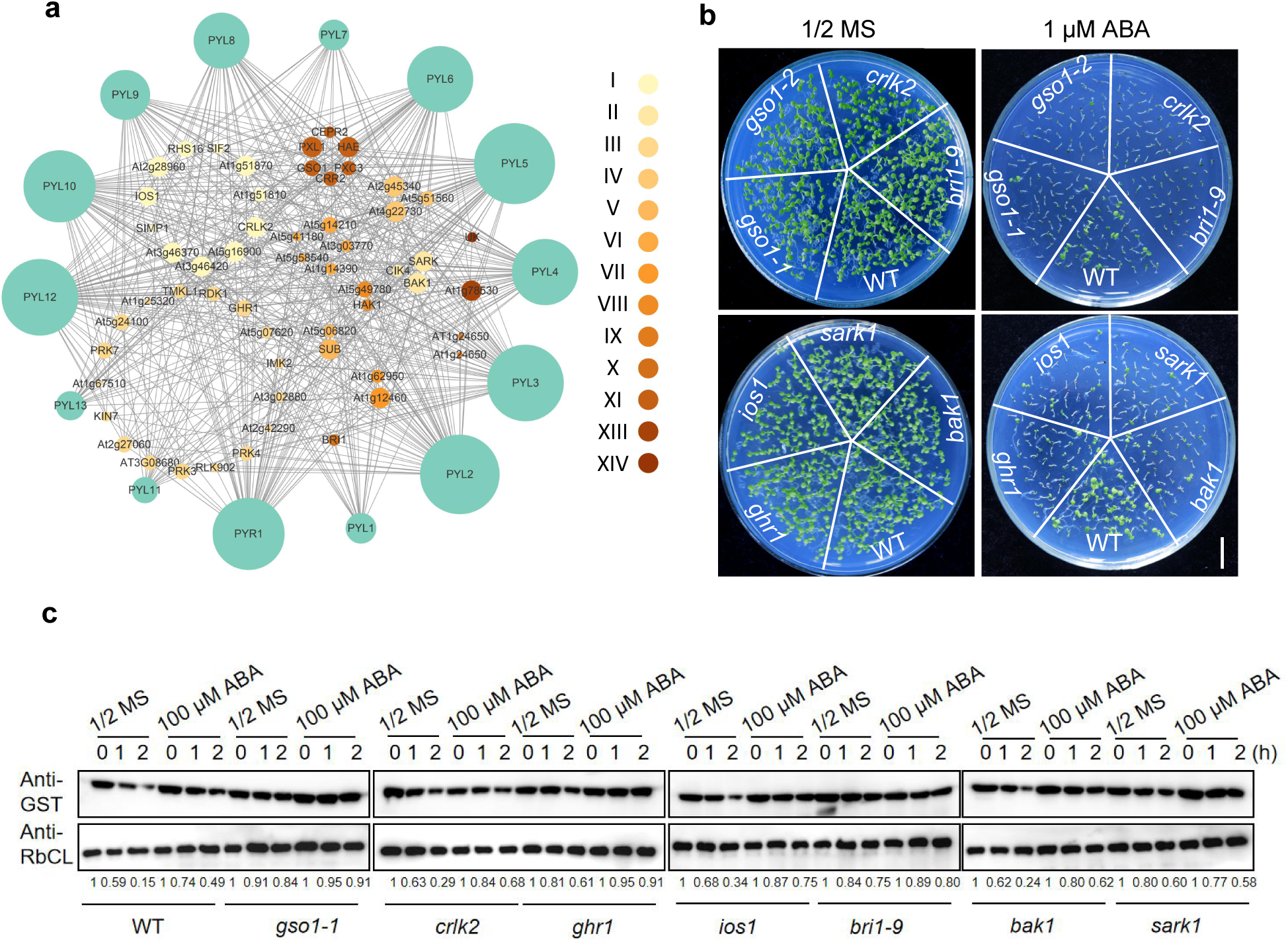
Membrane-based interactome of ABA receptors PYR1/PYLs with LRR-RLKs and the functional characterization of selected LRR-RLKs. **a,** Membrane yeast two-hybrid analysis revealing the interaction network between ABA receptors (PYR1/PYLs) and a panel of LRR-RLKs. The size of each circle is proportional to the number of its interacting partners, and the Roman numerals (I–XIV) represent the 13 LRR-RLK subfamilies. **b,** Germination phenotypes of wild-type (WT) and selected *lrr-rlk* mutant seeds grown for 10 days on 1/2 MS medium with or without 1 µM ABA. Scale Bar = 1 cm. **c,** *In vitro* degradation assay of recombinant PYL4-GST protein. PYL4-GST was incubated with soluble protein extracts prepared from WT or the indicated *lrr-rlk* mutant seedlings pretreated with or without 100 µM ABA for 1 or 2 h. Degradation was monitored by immunoblotting using an anti-GST antibody, with RbCL serving as the loading control.

To explore the functional relevance of these interactions, we selected seven candidate LRR-RLKs for further characterization. For each candidate, we obtained T-DNA insertion mutant lines and verified reduced transcript levels compared to the wild type (Supplementary Fig. 2a, b). In cotyledon greening assay, all mutant lines exhibited heightened sensitivity to ABA treatment relative to wild-type seedlings (Fig. 1b, Supplementary Fig. 2c), indicating that all seven LRR-RLKs act as negative regulators of ABA signaling. Given that each of the seven LRR-RLKs interacted with PYL4 in the yeast-based assays, we performed a cell-free degradation assay using recombinant purified PYL4 fused to glutathione *S*-transferase (PYL4-GST). When PYL4-GST was incubated with protein extracts from each mutant, we observed a consistently delayed in PYL4 degradation (Fig. 1c), implying a shared mechanism by which these LRR-RLKs regulates PYL stability. Together, these findings position the LRR-RLK network upstream of PYLs, modulating ABA sensitivity through post-translational control.

### GSO1 is a *bona fide* interacting partner and negative regulator of PYLs

Among the top candidate LRR-RLKs identified was GSO1. We confirmed its interaction with PYL1, PYL2, and PYL4 using bimolecular fluorescence complementation (BiFC) assays. Strong reconstituted yellow fluorescent protein (YFP) was detected when GSO1-nYFP (fused to the N-terminal half of YFP) was co-infiltrated with a PYL1-cYFP, PYL2-cYFP, or PYL4-cYFP (fused to the C-terminal half of YFP) in the leaves of *Nicotiana benthamiana* plants (Fig. 2a). In contrast, only weak YFP signal was observed when *GSO1-nYFP* was co-expressed with *PYL6-cYFP* or *PYL8-cYFP*, and no YFP signal was detected with *PYL5-cYFP*, *PYL10-cYFP*, or *PYL13-cYFP* (Supplementary Fig. 3). Luciferase complementation imaging (LCI) assays further corroborated these specific interactions (Fig. 2b–d). Additionally, GST pull-down assays using recombinant kinase domain of GSO1 (GSO1^KD^) and GST-tagged PYL1, PYL2, or PYL4 confirmed that GSO1^KD^-His was pulled down in each case (Fig. 2e–g). These results collectively demonstrate that GSO1 strongly interacts with PYL1, PYL2, and PYL4 both *in vivo* and *in vitro*.

**Figure 2.**
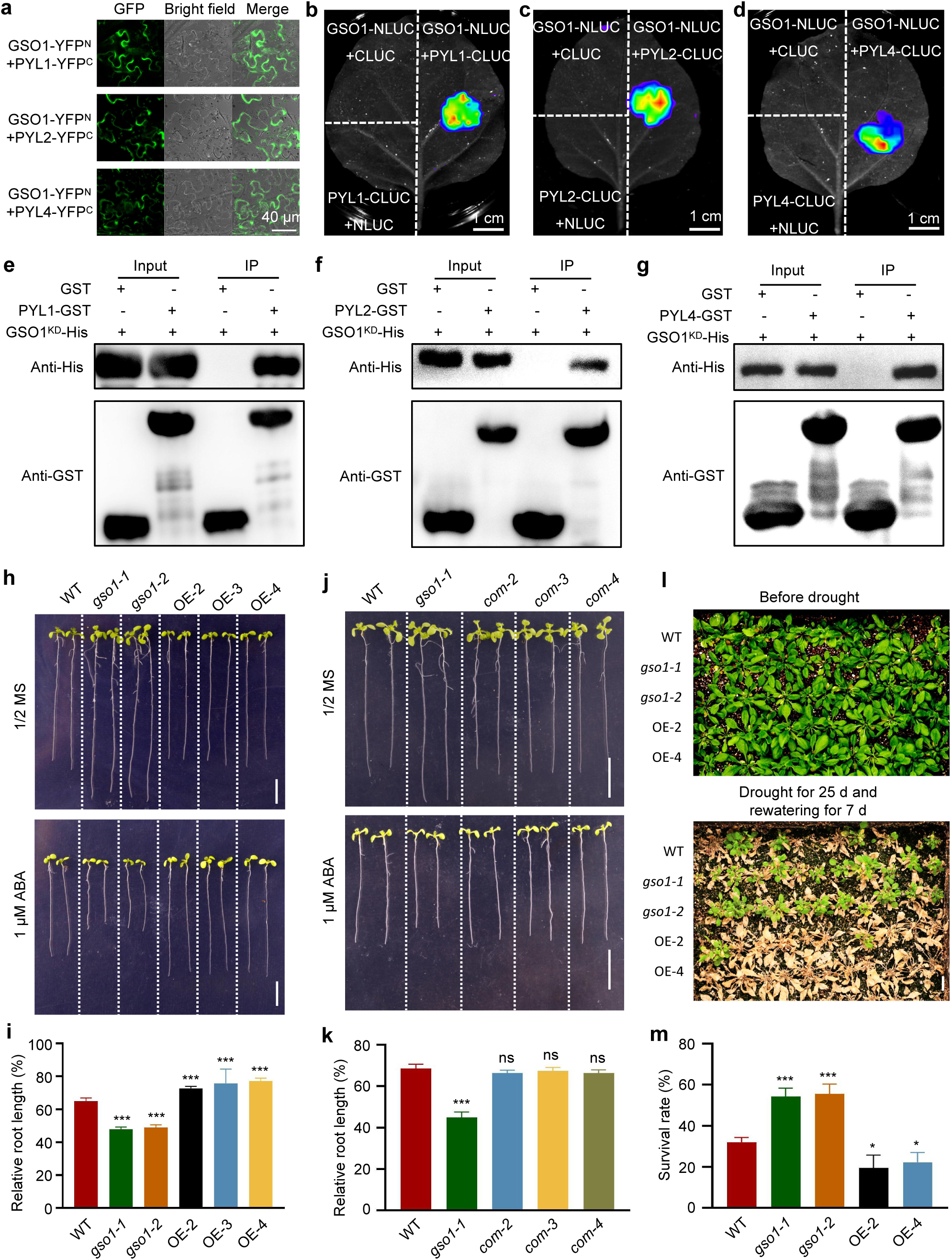
GSO1 physically interacts with PYR/PYL receptors and negatively regulates ABA and drought responses. **a,** Bimolecular fluorescence complementation (BiFC) assays showing the interaction between GSO1 and PYL1, PYL2, and PYL4 in *Nicotiana benthamiana* leaves. Scale bar: 40 μm. **b-d,** Firefly luciferase complementation imaging (LCI) assays confirming the interactions of GSO1 with PYL1, PYL2, and PYL4. Scale Bar: 1 cm. **e-g,** *In vitro* pull-down assays demonstrating direct biding of purified GSO1^KD^-His to PYL1-GST, PYL2-GST, or PYL4-GST, but not to GST alone. **h,** Phenotypes of 10-day-old WT, *gso1-1*, *gso1-2*, and GSO1-OE seedlings grown on 1/2 MS with or without 1 µM ABA. Scale bar: 1 cm. **i,** Quantification of relative root lengths from (**h**). Data are mean ± SEM (n = 20). Different lowercase letters indicate statistically significant differences (P < 0.05; one-way ANOVA). **j,** Genetic analysis of in ABA response. Phenotypes of WT, *gso1-1*, complementary (*com-2*, *com-3,* and *com-4*) lines grown for 10 days on on 1/2 MS with or without 1 µM ABA. Scale bar: 1 cm. **k,** Quantification of relative root lengths from (**j**). Data are mean ± SEM (n = 18). Different lowercase letters indicate statistically significant differences (P < 0.05; one-way ANOVA). **l,** Drought tolerance phenotypes of WT, *gso1-1*, *gso1-2*, and GSO1-OE plants. Three-week-old soil-grown plants were subjected to drought by withholding water for 25 days, the rewatered for 7 days to assess recovery. Scale bar: 2 cm. **m,** Survival rates of plants in (**l**), recorded 7 days after rewatering. Data are mean ± SEM (n = 3 biological replicates, each with at least 15 plants). Different lowercase letters indicate statistically significant differences (P < 0.05; one-way ANOVA).

To investigate the role of GSO1 in ABA signaling, we generated three independent *GSO1*-overexpression (*GSO1*-OE) transgenic lines; RT-PCR analysis indicated high accumulation of *GSO1* transcripts in these transgenic lines (Supplementary Fig. 4a). Under normal growth conditions on half-strength Murashige and Skoog (MS) medium, root length of two *gso1* mutant lines were longer than those of the wild type, whereas that of *GSO1*-OE lines was comparable to wild type. However, in the presence of 1 µM ABA, the *gso1* seedlings developed shorter roots and slower cotyledon expansion relative to the wild type, whereas *GSO1*-OE seedlings displayed longer roots and faster cotyledon expansion (Fig. 2h, i, Supplementary Fig. 4c, d). The short-root phenotype of *gso1* mutants under ABA treatment was fully rescued by introducing a genomic GSO1 construct under its native promoter (*proGSO1:GSO1*) (Fig. 2j, k). Taken together, these data establish that GSO1 acts as a negative regulator of ABA signaling.

Given that drought stress triggers ABA accumulation to promote plant acclimation, we investigated whether GSO1 is involved in drought response. After withholding water from 3-week-old plants for 25 days followed by 7 days of rewatering, both *gso1* mutants exhibited significantly enhanced drought tolerance and higher survival rates compared to wild-type plants (Fig. 3l, m). In contrast, *GSO1*-OE plants were slightly but significantly more susceptible to drought stress. These observations further support a negative regulatory role for GSO1 in *Arabidopsis* drought response.

**Figure 3.**
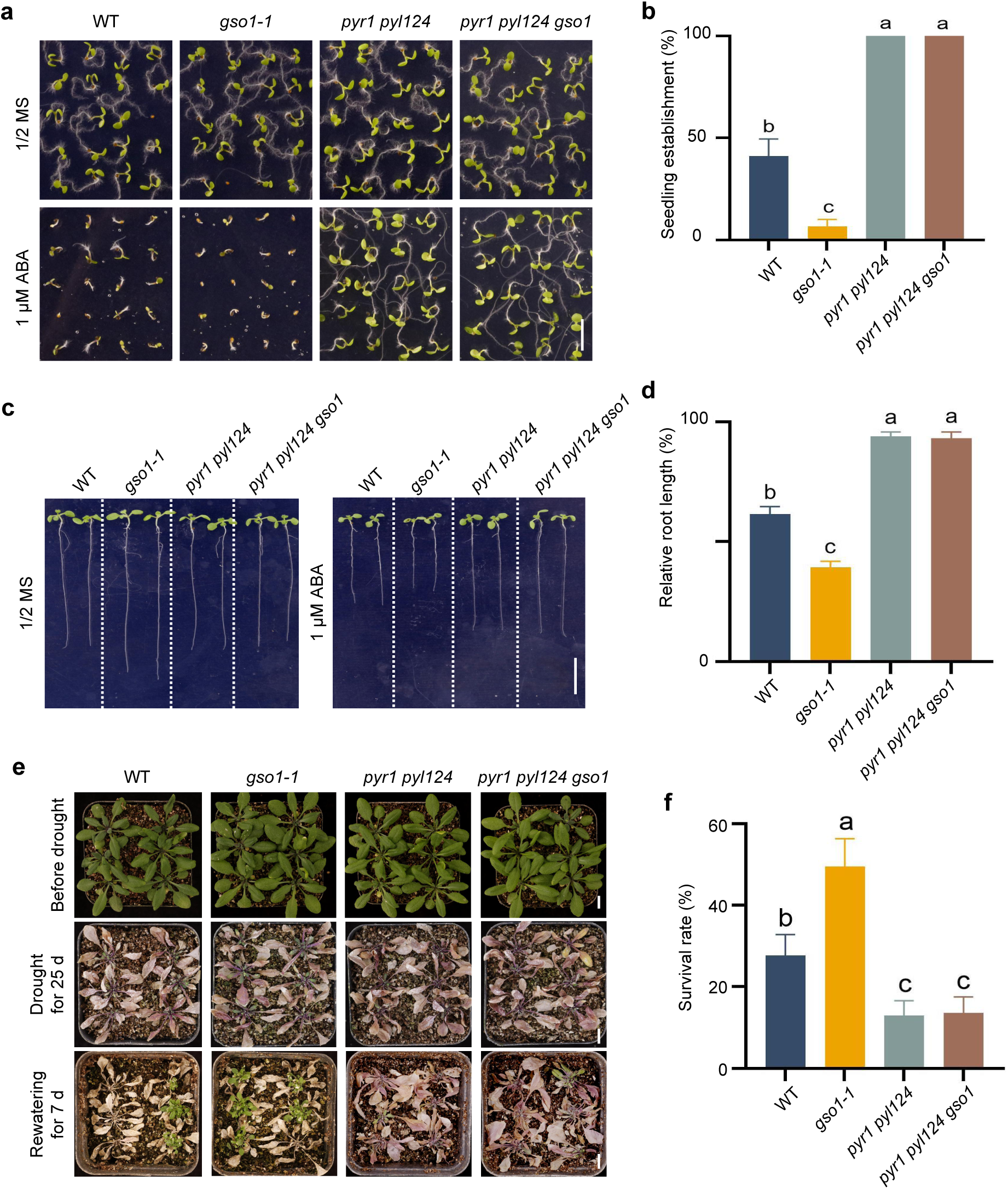
PYL receptors act downstream of GSO1 in regulating ABA responses. **a,** Phenotypes of 7-day-old WT, *gso1-1*, *pyr1 pyl124* ,and *pyr1 pyl124 gso1* seedlings grown on 1/2 MS medium with or without 1 μM ABA for 7 days. Scale bar: 0.5 cm. **b,** Quantification of seedling establishment rates from (**a**). Data are mean ± SEM (n = 3 biological replicates). Different lowercase letters indicate statistically significant differences (P < 0.05; one-way ANOVA). **c,** Phenotypes of 10-day-old WT, *gso1-1*, *pyr1 pyl124* ,and *pyr1 pyl124 gso1* seedlings grown on 1/2 MS medium with or without 1 μM ABA. Scale bar: 1 cm. **d,** Primary root lengths of seedlings from (**c**). Data are mean ± SEM (n = 18). Different lowercase letters indicate statistically significant differences (P < 0.05; one-way ANOVA). **e,** Drought tolerance of WT, *gso1-1*, *pyr1 pyl124* ,and *pyr1 pyl124/gso1* plants. Three-week-old soil-grown plants were deprived of water for 25 days, then re-watered for7-day to access recovery. Scale bar: 1 cm. **f,** Survival rates of plants in (**e**), recorded 7 days after rewatering. Data are mean ± SEM (n = 3 biological replicates, each with at least 18 plants). Different lowercase letters indicate statistically significant differences (P < 0.05; one-way ANOVA).

### GSO1 functions upstream of PYLs

To elucidate the genetic relationship between GSO1 and PYLs, we generated a *pyr1 pyl1 pyl2 pyl4 gso1* (*pyr1 pyl124 gso1* thereafter) quintuple mutant by hybridization and analyzed its ABA responses. The quintuple mutant exhibited strongly diminished sensitivity to ABA, with higher rates of cotyledon expansion and relatively longer primary roots than those of wild type. Notably, the ABA sensitivity of the *pyr1 pyl124 gso1* mutant was similar with that of the *pyr1 pyl124* quadruple mutant (Figs. 3a–d). The genetic interaction among PYLs and GSO1 was also evaluated under drought conditions. The *pyr1 pyl124 gso1* quintuple mutant displayed a similar drought sensitivity to that of the *pyr1 pyl124* quadruple mutant, but opposite from *gso1* on the bases of survival rate after water withholding and rewatering (Fig. 3e, f). These results demonstrate that GSO1 functions upstream of PYLs.

### GSO1 phosphorylates and promotes the degradation of PYL ABA receptors

Having established GSO1 as a functional kinase, we asked whether it directly phosphorylated PYLs. *In vitro* kinase assays using recombinant purified proteins revealed that GSO1^KD^-His phosphorylates PYL2-GST and PYL4-GST (Fig. 4a). To identify the specific phosphorylation sites, a liquid chromatography–tandem mass spectrometry (LC–MS/MS) analysis was performed using the recombinant purified PYL2-GST after *in vitro* phosphorylation by GSO1^KD^-His. Ser-33 and Ser-48 in PYL2, as well as a conserved Ser-65 in PYL4, were identified as putative phosphorylation sites (Supplementary Fig. 5a-c). Subsequent *in vitro* kinase assays with site-directed mutants demonstrated that Ser-48 in PYL2 and Ser-65 in PYL4 are the primary phosphorylation sites, as changing them to alanine (S48A and S65A, respectively) abolished GSO1-mediated phosphorylation (Fig. 4b).

**Figure 4.**
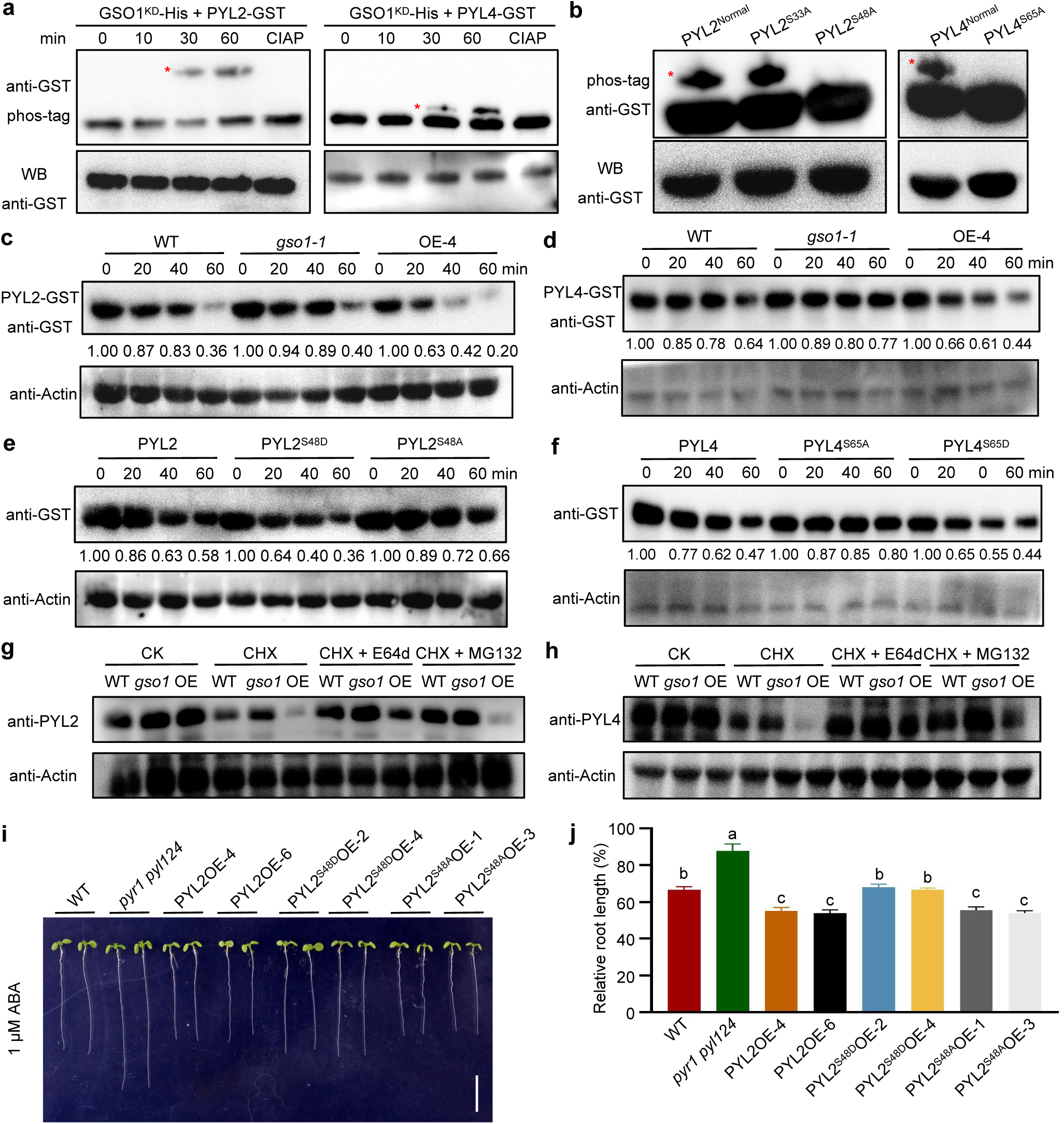
GSO1 promotes degradation of PYL2 and PYL4 through phosphorylation. **a,** *In vitro* kinase assays showing phosphorylation of PYL2 and PYL4 by GSO1, analyzed by Phos-tag SDS-PAGE. Treatment with calf intestinal alkaline phosphatase (CIAP) phosphorylation-dependent mobility shifts. **b,** *In vitro* kinase assays comparing phosphorylation of wild-type and phosphosite-mutant versions of PYL2 (PYL2-GST, PYL2^S33A^-GST, and PYL2^S48A^-GST) and PYL4 (PYL4-GST, PYL4^S65A^-GST) by GSO1. **c,d** *In vitro* degradation assays of PYL2-GST (**c**) and PYL4-GST (**d**) incubated with soluble protein extracts from 7-day-old WT, GSO1-OE-4, and *gso1-1* seedlings grown on 1/2 MS medium. Protein levels were detected by immunoblotting with anti-GST antibody. Actin serves as a loading control. **e,f** *In vitro* degradation assays of wild-type and phosphomimetic/mutant versions of PYL2 (PYL2, PYL2^S48A^, PYL2^S48D^; e) and PYL4 (PYL4, PYL4^S65A^, PYL4^S65D^; f) in soluble protein extracts from GSO1-OE-4 seedlings. Protein levels were monitored by anti-GST immunoblotting. **g,h** Degradation pathways analysis of PYL2 (g) and PYL4 (h) in 7-day-old WT, GSO1-OE-4, and *gso1-1* seedlings. Seedlings were treated for 4 h with the cycloheximide (CHX, 50 µM) alone, or together with MG132 (50 µM, proteasome inhibitor), E-64d (50 µM, vacuolar protease inhibitor), or both. Endogenous PYL2 and PYL4 protein levels were detected with specific antibodies. **i,** Phenotypes of 10-day-old WT, *pyr1 pyl124*, PYL2OE, PYL2^S48A^OE, and PYL2^S48D^OE seedlings grown on 1/2 MS medium supplemented with 1 μM ABA. Scale bar: 1 cm. **j,** Quantification of primary root lengths from (**i**). Data are mean ± SEM (n = 20). Different lowercase letters indicate statistically significant differences (P < 0.05; one-way ANOVA).

We next explored the functional consequences of this phosphorylation. Given that phosphorylation often regulates protein stability^31,32^, a cell-free degradation system was employed. Incubating recombinant purified PYL2-GST or PYL4-GST with plant extracts from *GSO1*-OE lines accelerated their degradation, whereas extracts from the *gso1-1* mutant stabilized them (Fig. 4c, d). Moreover, the phosphomimic variants PYL2^S48D^-GST and PYL4^S65D^-GST, in which the phosphorylated Ser residues were replaced with aspartic acid, were degraded more rapidly than their intact counterparts, whereas the nonphosphorylatable variants PYL2^S48A^-GST and PYL4^S65A^-GST were more stable (Fig. 4e, f). This degradation was suppressed by the addition of the proteasome inhibitor MG132 and the vacuolar degradation inhibitor E64d (Fig. 4g, h), suggesting the involvement of multiple turnover pathways.

To assess the physiological relevance of this phosphorylation *in planta*, transgenic *Arabidopsis* lines overexpressing intact *PYL2*, its nonphosphorylatable variant *PYL2^S48A^*, or its phosphomimic variant *PYL2^S48D^* were generated (Supplementary Fig. 6a–c). Under ABA treatment, *PYL2*-OE and *PYL2^S48A^*-OE lines were more sensitive to ABA than wild type, as evidenced by their pronounced inhibition of root growth. In stark contrast, the *PYL2^S48D^*-OE lines failed to enhance ABA sensitivity, exhibiting a response similar with wild-type seedlings (Fig. 4i, j, Supplementary Fig. 6d). This genetic evidence confirms that the GSO1-mediated phosphorylation of Ser-48 attenuates PYL2 function by promoting its degradation, thereby desensitizing seedlings to ABA.

### ABA and drought promote the degradation of GSO1

To further investigate the function of GSO1 in ABA signaling, the transgenic seedlings carrying a *proGSO1:GUS* reporter construct, consisting of the *GSO1* promoter driving the *β-Glucuronidase* (*GUS*) reporter gene, were we treated with ABA or polyethylene glycol (PEG) to mimic drought stress. Neither treatment affected the *GSO1* expression (Supplementary Fig. 7a). We then generated transgenic lines harboring a *35S:GSO1-GFP* transgene and treated them with ABA. GSO1 protein levels decreased rapidly within 3 hours and were nearly undetectable 24 hours after the onset of ABA treatment (Fig. 5a). A similar drop in GSO1 abundance under PEG treatment was observed (Supplementary Fig. 7b), indicating that ABA treatment and simulated drought stress promote the degradation of GSO1. To explore the underlying degradation mechanism, seedlings were treated with the endocytosis inhibitor E64d or the proteasome inhibitor MG132 in the presence of cycloheximide (CHX) and ABA. Treatment with E64d, but not MG132, markedly attenuated GSO1 degradation (Fig. 5b), suggesting that GSO1 is degraded via endocytosis.

**Figure 5.**
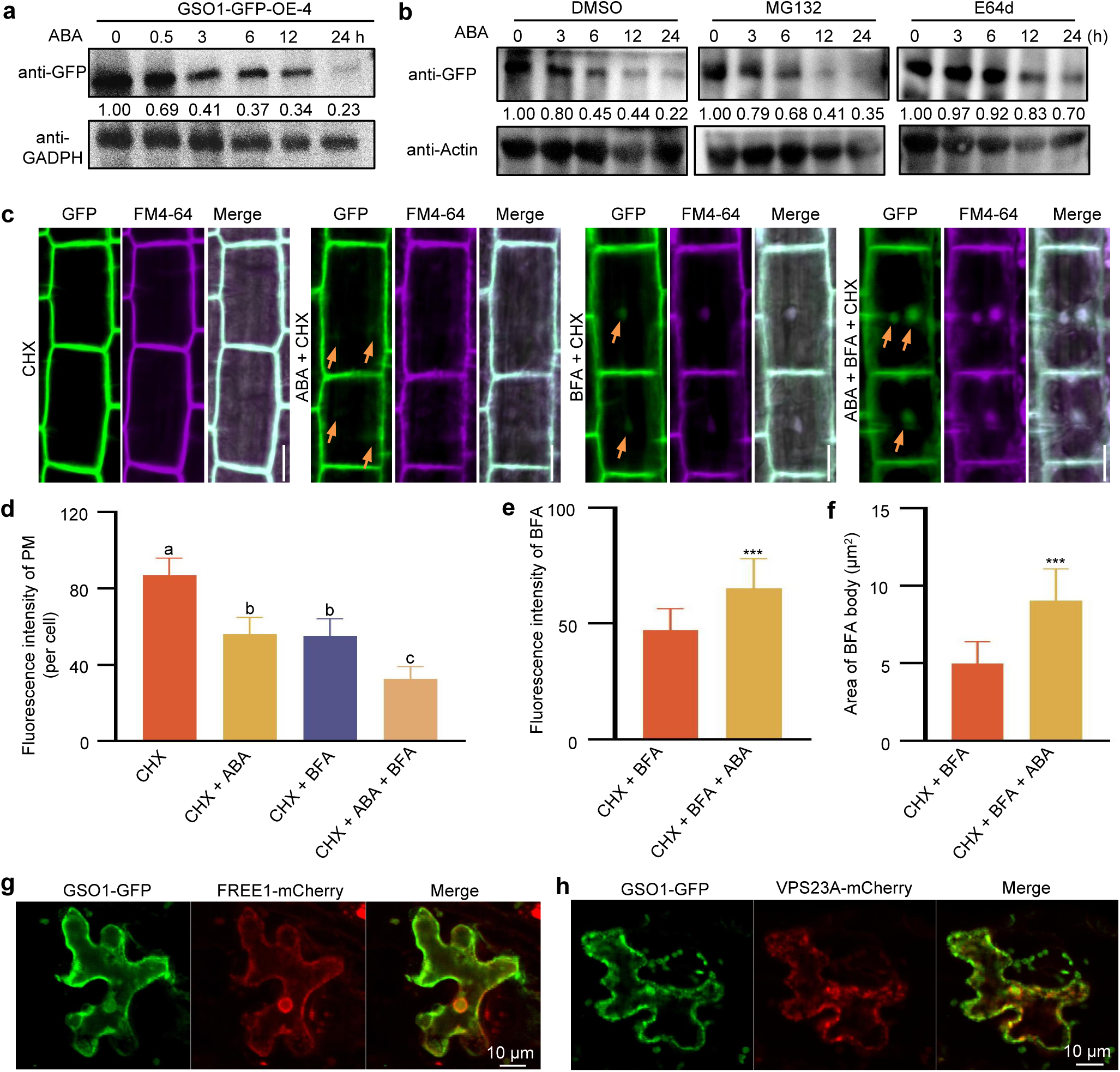
ABA promotes vacuolar degradation of GSO1. **a,** Time-course analysis of GSO1 protein stability following ABA treatment. Ten-day-old 35S:GSO1-GFP-OE-4 seedlings grown on 1/2 MS medium were treated with 50 μM ABA for the indicated durations (0 - 24 h). GSO1-GFP levels were detected by immunoblotting with an anti-GFP antibody; GADPH serves as the loading control. **b,** Effect of proteasomal and vacuolar inhibitors on ABA-induced GSO1 degradation. Seven-day-old 35S:GSO1-GFP seedlings were treated for 24 h with 100 µM ABA alone or together with 50 µM MG132 (proteasome inhibitor) or 50 µM E-64d (vacuolar protease inhibitor). GSO1-GFP abundance were assessed by anti-GFP antibody immunoblotting, with GADPH as loading control. **c,** Subcellular localization of GSO1-GFP under ABA and Brefeldin A (BFA) treatment. Seven-day-old 35S:GSO1-GFP seedlings were pre-treated with 50 µM cycloheximide (CHX) for 1 h, then co-treatment for 0.5 h with CHX (50 µM) plus one of the following: ABA (100 µM) plus FM4-64 (2 µM); BFA (50 µM) plus FM4-64 (2 µM); or BFA (50 µM) plus ABA (100 µM) plus FM4-64 (2 µM). Roots were imaged by confocal microscopy. Yellow arrowheads indicate BFA compartments containing GSO1-GFP. Scale bar:10 µm. Each treatment was analyzed in >15 images. **d-f,** Quantitative analysis of plasma-membrane fluorescence intensity (**d**), fluorescence intensity of BFA compartments (**e**), and area of BFA compartments (**f**) from (**c**). Data are mean ± SD (n = 20). Statistical significance was assessed using two-tailed unpaired *t*-test (*P < 0.05, **P < 0.01, ***P < 0.001). **g,h** Colocalization of GSO1 with the ESCRT components VPS23A and FREE1. GSO1-GFP was co-expressed with VPS23A-mCherry (**g**) or FREE1-mCherry (**h**) in *N. benthamiana* leaves. Tissues were treated with 50 µM ABA for 0.5 h before simultaneous detection of GFP and mCherry signals. Scale bar: 10 µm. All experiments were independently repeated at least three times with consistent results.

We also examined the effect of ABA treatment on GSO1 subcellular localization. GSO1-GFP was located at the plasma membrane with CHX treatment, whereas the addition of ABA prompted GSO1-GFP internalization into punctate structures. Co-localization studies using the transport inhibitor brefeldin A (BFA) confirmed these punctate as endocytic compartments. Combined treatment with CHX, BFA, and ABA produced more numerous and intensely fluorescent BFA bodies than CHX and BFA (Fig. 5c–f), providing direct evidence that ABA enhances GSO1 endocytosis for vacuolar delivery.

Since vacuolar degradation involves the endosomal sorting complex required for transport (ESCRT)^33–35^, we tested whether GSO1 colocalizes with core ESCRT-I components. *GSO1-GFP* with a *FREE1-mCherry* construct encoding a fusion between FYVE DOMAIN PROTEIN REQUIRED FOR ENDOSOMAL SORTING 1 (FREE1) and the red fluorescent protein mCherry or a *VACUOLAR PROTEIN SORTING 23A* (*VPS23A*)-*mCherry* construct were co-expressed in the leaves of *N. benthamiana* plants, followed by treatment with ABA. FREE1-mCherry and VPS23A-mCherry both appeared as punctate structures in the cytoplasm, and GSO1-GFP strongly colocalized with these markers (Figs. 5g, h). These analyses confirm that ABA promotes GSO1 degradation through the endocytic-vacuolar pathway. Thus, under drought stress, ABA accumulation accelerates the removal of GSO1, thereby relieving its inhibition on PYL receptors and amplifying ABA signaling.

### The CIF–GSO1 module is inhibited by ABA and drought stress

The CIF peptides, which are GSO1 ligands, act upstream of this pathway^26,29^. Drought stress and ABA treatment significantly downregulated the expression levels of *CIF1* and *CIF2* (Fig. 6a, b). However, when fluridone (Flu), an ABA biosynthesis inhibitor, was applied before drought stress, the repression of *CIF1* and *CIF2* imposed by PEG treatment disappeared (Fig. 6c). Paralleling this finding, the ABA-induced suppression of *CIF1* and *CIF2* expression was also abolished in *pyr1 pyl124* seedlings (Fig. 6d), providing genetic evidence that this repression acts through the canonical PYL receptor pathway.

**Figure 6.**
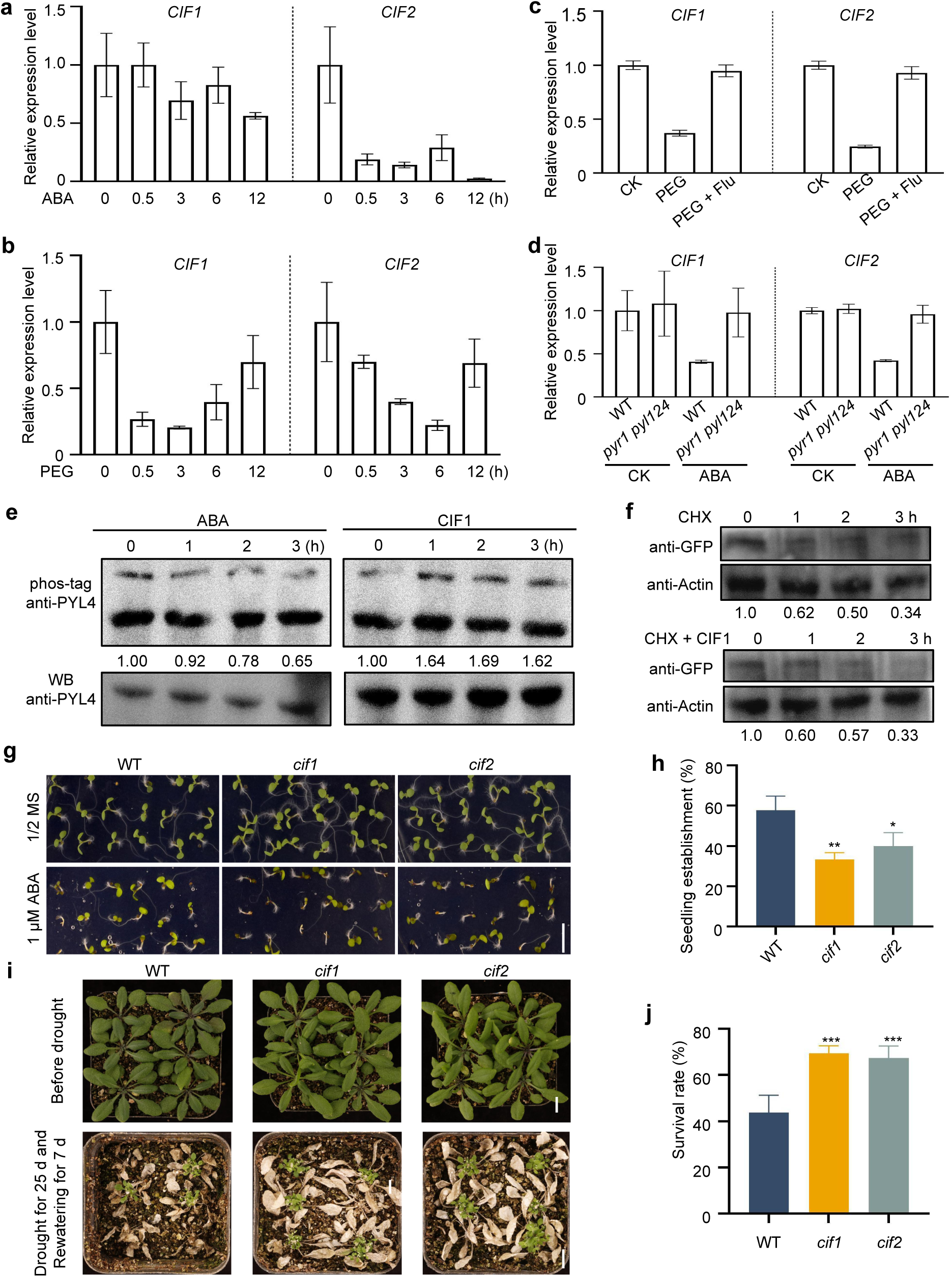
The CIF-GSO1 module is suppressed by drought stress. **a,b,** Transcript levels of *CIF1* and *CIF2* in 10-day-old wild-type seedlings treated with 100 μM ABA or 30%PEG (to simulate drought stress) for indicated durations. Expression was normalized to *18S* rRNA. Data are ± SEM (n = 3 biological replicates). Different lowercase letters indicated statistically significant differences (P < 0.05; one-way ANOVA). **c,** *CIF1* and *CIF2* expression in 7-day-old wild-type seedlings under control, 30%PEG, or 30%PEG plus 50 μM fluridone (ABA biosynthesis inhibitor). Seedlings were pre-treated with 50 μM fluridone for 6 h. Transcript levels were normalized to *18S* rRNA. Data ate means ± SEMs (n = 3). Different lowercase letters indicate statistically significant difference (P < 0.05; one-way ANOVA). **d,** *CIF1* and *CIF2* transcript levels in 7-day-old WT and *pyr1 pyl124* mutant seedlings with or without 100 μM ABA treatment. Expression was normalized to *18S* rRNA. Data are means ± SEM (n = 3). Different lowercase letters indicate statistically significant difference (P < 0.05; one-way ANOVA). **e,** *In vitro* kinase assay of GSO1 activity toward PYL4 in the presence of 100 μM ABA or 100 nM CIF1. Reactions were stopped at the indicated times, and PYL4 phosphorylation was analyzed by Phos-tag SDS-PAGE. **f,** Analysis of GSO1 protein stability. Ten-day-old 35S:GSO1-GFP-OE-4 seedlings were treated with 50 μM CHX alone or together with 100 nM CIF1 for the indicated times. GSO1-GFP protein levels were detected by anti-GFP antibody immunoblotting. Actin serves as loading control. **g,** Germination phenotypes of WT, *cif1*, and *cif2* mutants grown for 10 days on 1/2 MS medium with or without 1 µM ABA. Scale bar: 0.5 cm. **h,** Quantification of seedling establishment rates from (**g**). Data are mean ± SEM (n = 3). Different lowercase letters indicate statistically significant differences (P < 0.05; one-way ANOVA). **i,** Drought tolerance of WT, *cif1*, and *cif2* plants. Three-week-old soil-grown plants were deprived of water for 25 days, then re-watered for 7 days to assess recovery. Scale bar: 2 cm. **j,** Survival rates of plants in (**i**). Data are mean ± SEM (n = 3 biological replicates, each with at least 18 plants). Different lowercase letters indicate statistically significant differences (P < 0.05; one-way ANOVA).

To determine how CIF peptide levels directly influence the kinase activity of GSO1, *in vivo* kinase assays using protein extracts from *GSO1*-OE seedlings were performed. The addition of a synthetic CIF1 peptide enhanced GSO1 kinase activity towards its substrate PYL4, whereas ABA treatment had the opposite effect (Fig. 6e). However, in contrast to its potent effect on GSO1 kinase activity, CIF1 treatment did not affect the stability of GSO1-GFP in CHX chase assays (Fig. 6f). Phenotypically, *cif1* and *cif2* T-DNA insertion mutants were more sensitive to ABA treatment in cotyledon greening assays and more resistant to drought stress than the wild type (Fig. 6g–j; Supplementary Fig. 8). These findings establish a direct regulatory link: under normal non-stress conditions, CIF peptides activate GSO1 to phosphorylate and degrade PYLs. During drought stress, suppressed *CIF* expression leads to lower CIF peptide abundance and GSO1 activity, thereby releasing the brake on PYL stability and ABA signaling.

## Discussion

The LRR-RLK family, comprising 225 members in *Arabidopsis* alone, is indispensable for a multitude of plant functions, from development and phytohormone signaling to innate immunity^36^. While these studies highlight the specificity of individual LRR-RLKs, they also underscore a fundamental gap in our understanding. An emergent property of these receptors, their capacity to form higher-order networks for integrated signal regulation and developmental coordination, is an exciting and open research frontier.

Drought stress imposes a major constraint on global agricultural productivity^37^. Plants have evolved intricate signaling networks that mitigate the effects of water deficit, with the ABA pathway serving as a central regulator^38,39^. Although the abundance of ABA receptors is a critical determinant of signaling output, the upstream mechanisms that dynamically control PYL stability remain poorly understood. Our study fundamentally redefines the upstream architecture of ABA signaling, shifting from a linear perspective to a network model in which the abundance of PYL receptors is regulated by an extensive suite of LRR-RLKs. The finding that 53 out of 196 tested LRR-RLKs physically associate with PYR1 and/or PYLs (Supplementary Fig. 1), together with the finding that loss of function of seven distinct members consistently enhanced PYL4 stability and ABA sensitivity (Fig. 1b, c), indicates that control of receptor availability constitutes a key mechanism for signal integration. Our results position PYR1 and PYLs at a critical signaling nexus, enabling them to assimilate inputs from a broad set of signals, spanning phytohormones, pathogens, symbionts, and developmental cues perceived by different LRR-RLKs^14^, to compute context-appropriate ABA outputs. This framework also offers a mechanistic basis for the crosstalk between ABA and other signaling pathways.

Within this network, our in-depth analysis of the CIF–GSO1 module reveals a regulatory principle that is elegant in its design and profound in implication. We demonstrate that under non-stress conditions, GSO1 acts as a constitutive brake on PYL stability by phosphorylating PYL2 at Ser-48 and PYL4 at Ser-65, modifications that promote PYL degradation and attenuate ABA signaling to favor growth (Fig. 4). Under drought stress, however, the ABA-mediated suppression of *CIF* expression alleviates this constraint imposed on GSO1 (Fig. 6), constituting a derepression mechanism that rapidly sensitizes cells to ABA and facilitates the transition from growth to defense. Furthermore, ABA treatment and drought stress promote GSO1 endocytosis and subsequent vacuolar degradation (Fig. 5). Through its dual role in growth and ABA signaling, GSO1 establishes a direct molecular link between developmental programs and environmental acclimation, thereby functioning as a central integrator governing resource allocation between soil exploration and stress avoidance.

The conceptual significance of this mechanism is further illuminated by comparison with the CEPR2-mediated pathway. Although both pathways converge on PYL stabilization to potentiate ABA signaling, they operate through opposing regulatory logics. The CIF–GSO1 module functions via derepression: drought and ABA suppress *CIF* expression, thereby inactivating GSO1 and releasing the brake on PYL accumulation. By contrast, the CEP–CEPR2 pathway operates through ligand-inhibited feedback, wherein drought-induced CEPs inhibit CEPR2 kinase activity, likewise leading to PYL stabilization^12,13^. The coexistence of these two pathways, both enhancing ABA perception via opposing regulatory strategies, creates a tunable and robust system for modulating stress response dynamics, offering a sophisticated biochemical solution to the growth–defense trade-off.

These findings paint a picture of remarkable regulatory sophistication, with distinct kinases targeting different residues for phosphorylation. Indeed, GSO1 phosphorylates PYL4 at Ser-65 (Fig. 4), CEPR2 phosphorylates PYL4 at Ser-54^12^, ARABIDOPSIS EL1-LIKE (AELs) phosphorylates PYL1 at Ser-136 and Ser-183^40^, and CALCINEURIN B-LIKE-INTERACTING PROTEIN KINASE 1 (CIPK1) phosphorylates PYL4 at Ser-129^40,41^. The observation that distinct kinases phosphorylate different residues within PYLs raises the possibility of a “phospho-code”, wherein combinatorial phosphorylation patterns direct specific outcomes, such as preferential degradation via the proteasome or vacuole, or altered interactions with PP2Cs or other partners. Such specificity, combined with divergent ligand-regulation logics, endows the network with the potential to generate a broad spectrum of dynamic responses from a limited set of components.

Looking ahead, a major challenge lies in deciphering the architecture and dynamics of this network. Key questions remain: How are specific LRR-RLK partners recruited to PYLs? Which is the specific phosphorylation site for a specific LRR-RLK? How are their activities coordinated temporally and spatially to process conflicting signals? And which E3 ubiquitin ligases and associated factors execute the final degradation steps of LRR-RLKs? Addressing these questions will be essential for a holistic understanding of how plants dynamically modulate ABA sensitivity.

In summary, we have uncovered a regulatory network that confers robustness, flexibility, and integrative capacity to ABA signaling. Focusing on GSO1 and CEPR2, we propose a “dual-safeguard” feedback model in which the CIF–GSO1–PYL and CEP–CEPR2–PYL modules act in concert to balance growth and drought resilience (Fig. 7). Under normal conditions, high CIF levels sustain GSO1 activity, whereas low CEP levels maintain CEPR2 activity, with both activated LRR-RLKs promoting PYL degradation and suppressing ABA signaling to favor growth. Under drought stress conditions, however, ABA accumulation triggers two complementary responses: suppression of *CIF* expression and vacuolar degradation of GSO1, both of which inactivate the GSO1 brake, whereas induction of *CEP* expression inhibits CEPR2 kinase activity. Together, these mechanisms ensure rapid PYL accumulation, robust ABA pathway activation, and efficient drought response. Once stress conditions recede, the system resets. Thus, GSO1 and CEPR2 function as central molecular switches that fine-tune the balance between growth and stress response.

**Figure 7.**
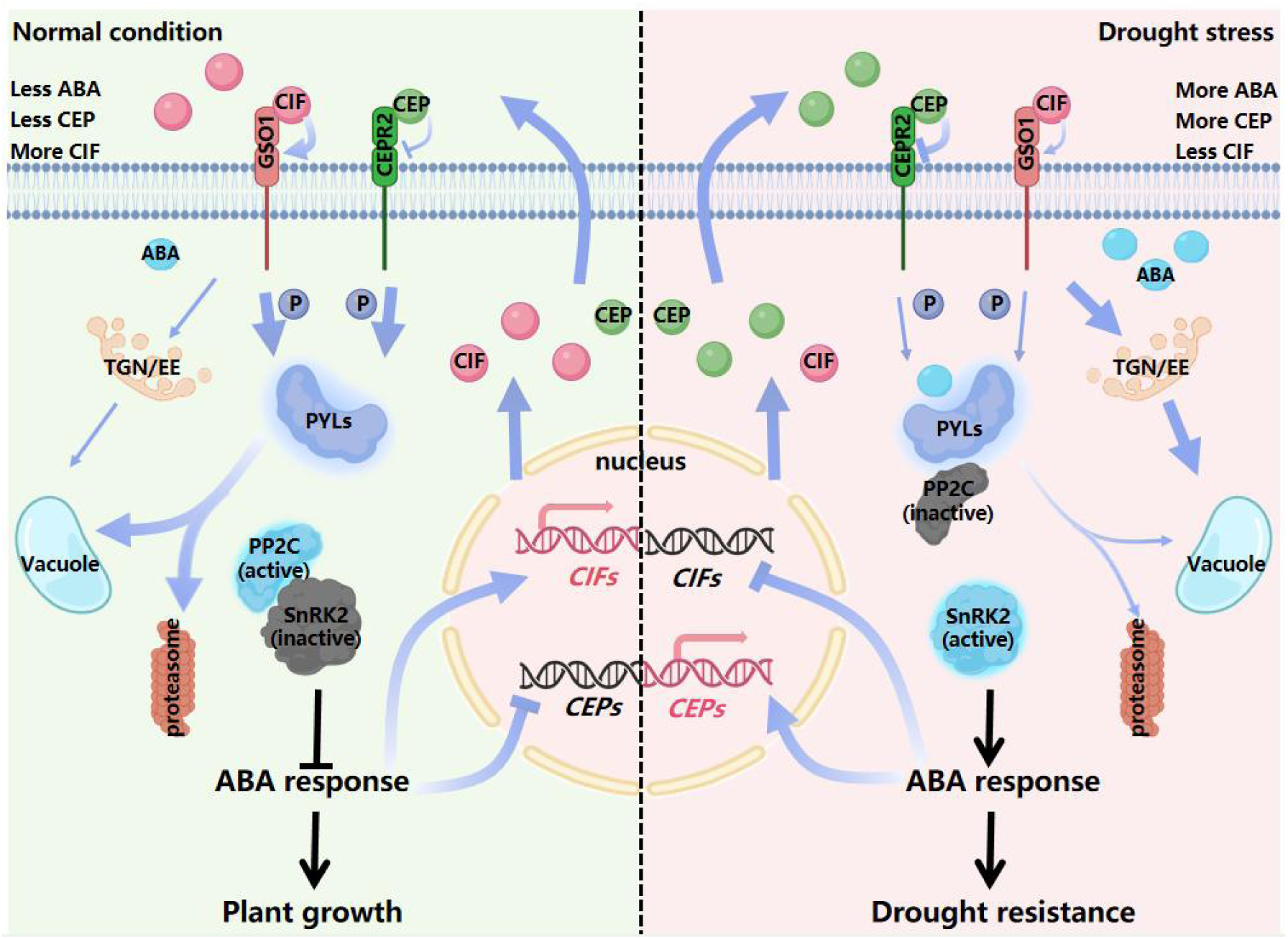
Model for the coordinated interaction between the CIF-GSO1-PYL and CEP-CEPR2-PYL modules in balancing growth and drought resistance. Under normal conditions, low ABA signaling permits relatively high expression of *CIFs* but low expression of *CEPs*. Active GSO1 and CEPR2 kinases phosphorylate PYL receptors, targeting them for degradation via the 26S proteasome and the vacuole. This maintains low ABA signaling and promotes growth, completing a homeostatic circuit. During drought, accumulated ABA initiates a positive feedback loop: it represses *CIFs* and induces *CEPs,* thereby inactivating their corresponding kinases, and concurrently promotes GSO1 degradation. These changes stabilizes PYL proteins, significantly enhancing ABA perception and signal output. The potentiated ABA signaling further reinforces *CIFs* suppression and *CEPs* induction, closing the circuit to lock thesystem into a high-sensitivity state, ensuring robust drought resistance. In the schematic, arrows indicate regulation (activation or promotion), and blunt-ended lines denote negative regulation (inhibition). Line thickness reflects relative signal strength. Figure created with BioRender.com.

## Methods

### Plant materials and growth conditions

Wild type (WT) *Arabidopsis thaliana* (L.) Heynh. cv. ‘Columbia’ was used as control. T-DNA insertion lines (e.g., SALK_137388 for *ios1*, SALK_111290 for *sark1*, SALK_034523 for *bak1*, SALK_031493 for *ghr1*, SALK_064029 for *gso1-1*, SALK_034282 for *gso1-2*, and *bri1-9*) were provided by Dr. Chunming Liu and Xiufen Song (Key Laboratory of Plant Molecular Physiology, Institute of Botany, Chinese Academy of Sciences, Beijing, China). Seedlings of these lines were treated on half-strength (1/2) Murashige and Skoog (MS) medium (1.5% sucrose and 0.85% agar) with or without ABA. GSO1, PYL2 and PYL4-overexpressing lines were generated by introducing their coding regions into the pCAMBIAsuper1300 vector (Liu et al., 2020)^42^. The transformation of *Arabidopsis* was performed using the floral-dip method with *Agrobacterium tumefaciens* strain GV3101. T3 transgenic plants were selected on half-strength MS medium supplemented with 50 mg L^−1^ kanamycin or 25 mg L^−1^ hygromycin B and confirmed by RT-qPCR using the primers listed in Supplemental Table1.

The homozygous single and quintuple mutants (*gso1-1*, *gso1-2*, and *pyr1 pyl124 gso1*) were verified using RT–PCR with the primers listed in Supplemental Table1. All selected *Arabidopsis* plants were grown in a greenhouse under a 16-h light/8-h dark cycle at 23℃ with a light intensity of 9600 lux.

### MbSUS assay

The MbSUS assay was performed as described previously; detailed descriptions of associated experimental principles and methods are presented in Yu et al^12^. In brief, for Nub fusions, PCR products were cloned and transformed with pNXgate, cleaved with *EcoRI* and *SmaI*, and transformed into the yeast strain THY.AP5. Strains were selected on synthetic dropout (SD) medium lacking tryptophan (W) and uracil (U). For Cub fusions, PCR products were cloned and transformed with pMetYCgate, cleaved with *Pst*I/*Hind*III, and transformed (along with PCR products) into the yeast strain THY.AP4. Transformants were further selected on SD medium lacking leucine (L). Clones from each transformant were incubated on SD medium lacking leucine, tryptophan, and uracil (SD-Trp/-Leu/-Ura; SD-WLU) at 30°C for 3 days. To detect protein interactions, we spotted colonies onto control medium (SD-WLU) and onto selection medium lacking leucine, tryptophan, uracil, adenine, and histidine (SD-Trp/-Leu/-Ura/-Ade/-His; SD-WLUAH). Colony growth was monitored for 3–6 days.

### MIND construction

MIND was inferred through the method described previously^43^, and obtained from the STRING databases (https://cn.string-db.org/, version 12.0 accessed on 7 April 2025)^44^. The sequence relationship data of *Arabidopsis* was downloaded from STRING with medium confidence, respectively. The network was visualized using Cytoscape software v 3.10.3 (http://www.cytoscape.org accessed on 5 December 2024)^45^.

### Measurements of relative germination rate, green cotyledon rate, and root length

Measurement of relative germination rate was performed as previously described^46^. Briefly, seeds were surface-sterilized and then incubated for three days on the 1/2 MS medium with or without 1 μM ABA at 4°C and in the dark at 4°C for three days. Germination rates of the WT or the mutant lines were determined at two days post incubation. A germinated seed was defined as the emergence of radicle through seed coat. The rate of plants with green cotyledons per line was determined after 3 days. For growth analysis, the sterilized seeds were incubated on the 1/2 MS medium with or without 1 μM ABA for ten days. Primary root of each seedling was measured using a ruler at 10 days post incubation. Experiment was repeated 3 times (n=20).

### LCI and BiFC assays

LCI assays were performed as previously described^47^. In brief, the full-length coding sequence (CDS) of GSO1 was cloned into the pCAMBIA1300-nLUC (nLUC) vector to generate the GSO1-nLUC construct, and the full-length CDS of PYLs was cloned into the pCAMBIA1300-cLUC (cLUC) vector to generate the PYLs-cLUC construct. Next, 1 ml samples of Agrobacterium tumefaciens cells harboring GSO1-nLUC, PYLs-cLUC, or nLUC were mixed to obtain the following combinations, each with a final optical density at 600 nm (OD600) of 1.0: GSO1-nLUC + cLUC, PYLs-cLUC + nLUC, and GSO1-nLUC + PYLs-cLUC. Each combination of *A. tumefaciens* cells was infiltrated separately into *N. benthamiana* leaves. *N. benthamiana* plants were grown at 26°C for 60 h. Five minutes before detection, 0.2 mM luciferin (Promega, Madison, WI, USA) was sprayed onto the treated leaves, and luciferase activity was measured using a cooled charge-coupled device (Lumina II system; PerkinElmer, Waltham, MA, USA).

BiFC assays were performed as described previously^48^. In brief, the full-length CDS of GSO1 (without stop codon) was cloned into the pSPYNE-35S vector to generate the GSO1-YFPN construct, and the full-length CDS of PYLs (without stop codon) was cloned into the pSPYCE-35S vector to obtain the PYLs-YFPC construct. A. tumefaciens cells harboring GSO1-YFPN and PYLs-YFPC were mixed with 10 ml of MMA medium (10 mM MgCl_2_, 50 mM MES [2-(N-morpholino) ethanesulfonic acid], and 20 mM acetosyringone [pH 5.6]) to yield a final OD600 of 1.0. The cell mixtures were injected into *N. benthamiana* leaves by gently pressing a disposable syringe to the abaxial surface of a fully expanded leaf (approximate width: 3 cm at the midpoint). Plants were grown at 26°C for 36–60 h, and YFP signals in the leaves were detected at 488 nm using an LSM 51 confocal laser scanning microscope (Zeiss, Germany).

### Pull-down assays

To characterize the interactions among GSO1, PYL1, PYL2 and PYL4, the CDS of PYLs was fused with GST in the pGEX-4T-3 vector to generate pGEX-4T-3-GST-PYLs (PYLs-GST); and the kinase domain (899–1249 aa) of GSO1 was fused with the His tag in the pET30a-His vector to generate pET30a-GSO1^KD^-His (GSO1^KD^-His). The resultant PYLs-GST and GSO1^KD^-His constructs were transformed into competent *E. coli Rosetta* cells. The transformed cells were cultured in 500 ml of Luria-Bertani medium at 37°C to an OD600 of 1.0, and then induced with 0.8 mM isopropyl b-D-thiogalactoside (IPTG) for 12 h at 16°C for protein expression. Next, *E. coli* cells were obtained by centrifugation at 6000 rpm for 5 min at 4°C. The pellet was resuspended in 5 ml of ddH_2_O. Lysates were obtained by ultrasonication (JY92-II; Scientz Biotechnology, Ningbo, China) with the following parameters: operating power, 300 W; working time, 10 s; interval time, 5 s; cycles, 30. Lysates were clarified by centrifugation at 8000 rpm for 10 min at 4°C. The GSO1^KD^-His protein was purified using the His-Tagged Protein Purification Kit (CWBIO, Beijing, China), and PYL2-GST and PYL4-GST were purified using Pierce Glutathione Spin Columns (Thermo Fisher Scientific, Waltham, MA, USA). In the pull-down assay, each 50 mg of PYLs-GST and GSO1^KD^-His were incubated in 1 ml of binding buffer (50 mM Tris–HCl and 150 mM NaCl [pH 8.0]) at 4°C for 2 h with constant slight shaking. After incubation, GST proteins were purified with Pierce Glutathione Spin Columns, eluted, and analyzed using anti-His antibodies (CWBIO). The primers used in this experiment are listed in Supplemental Table1.

### Phenotype analysis

Plants of different genotypes were grown under the same conditions in a controlled growth chamber. Seeds were collected at the same time. For drought treatment, plants grown on 1/2 MS medium for 7 days were transplanted into soil for about 2 weeks with sufficient watering, followed by 12 days of drought stress (withholding irrigation). In each experiment, six plants were grown in a small pot, or different plants were cultivated in a large pot with 12 individuals of each species spaced apart. The plants were grown under a photoperiod of 16 hours light/8 hours darkness at 22°C; at least three independent experiments were performed. Photos were taken after 25 days of drought treatment followed by 7 days of rewatering. The survival rates were then calculated.

### RNA extraction, RT–PCR, and qRT–PCR

Total RNA was extracted from 7-day-old seedlings grown on 1/2 MS medium with or without 1 μ M ABA using TRIzol reagent (Invitrogen, Carlsbad, CA, USA) or Universal Plant Total RNA Extraction Kits (Spin-column)-I (BioTeke, Beijing, China). The cDNA used for quantitative RT–PCR (qRT–PCR) was synthesized using Evo M-MLV RT Kit with gDNA Clean for qPCRII AG11711(Accurate Biotechnology [Hunan] Co., Ltd). qRT–PCR was performed using the ChamQ Universal SYBR qPCR Master Mix (Vazyme Biotech) on a CFX96 instrument (Bio-Rad, Hercules, CA, USA). UBC21 and UBQ10 were used as internal controls for qRT–PCR. The cDNA used for RT–PCR was synthesized using the PrimeScript 1^st^ strand cDNA Synthesis Kit (Takara, Osaka, Japan). RT–PCR cycling conditions were as follows: denaturation at 94°C for 5 min, 23–34 cycles of amplification, and final elongation at 72°C for 5 min. *EF-1a* was used as the internal control for RT–PCR. The primers used are listed in Supplemental Table1.

### *In vitro* kinase assays

To investigate possible sites at which GSO1 phosphorylates PYL2 and PYL4, we carried out *in vitro* kinase assays. CDSs encoding the mutant protein at Ser33 and Ser 48 of PYL2 and Ser 65 of PYL4 were mutated to alanine (A) or aspartate (D) and were fused with GST tag in the pGEX-4T-1 vector to generate pGEX-4T-1-GST-PYL2S33A/D (PYL2S33A/D-GST), pGEX-4T-1-GST-PYL2S48A/D (PYL2S48A/D-GST), and pGEX-4T-1-GST-PYL4S65A/D (PYL4S65A/D-GST). Transformation, induction, and purification of these fusion proteins were performed as described above. GSO1^KD^-His was purified using affinity chromatography. *In vitro* kinase assays were carried out using purified GSO1^KD^-His and PYL2-GST, PYL2S33A/D-GST, PYL2S48A/D-GST, PYL4-GST or PYL4S65A/D-GST. Fifty micrograms of PYL2-GST, PYL2S33A/D-GST, PYL2S48A/D-GST, PYL4-GST or PYL4S65A/D-GST were combined with 0.5 mg of GSO1^KD^-His in 50 ml of reaction buffer (25 mM HEPES [pH 7.2], 1 mM DTT, 50 mM NaCl, 2 mM EGTA, 5 mM MgSO_4_, and 50 mM ATP). The reaction mixtures were incubated at 30°C for 0-60 min, and the reaction was terminated by the addition of loading buffer. Unincubated reaction mixture (at 0 min) was used as the negative control. Proteins were then fractionated by SDS–PAGE and Mn^2+^-Phos-tag–PAGE (50 mM Phos-tag and 100 mM Mn^2+^). After a 30 min incubation at 37°C, 0.01 U/ml CIAP (Promega, Madison, WI, USA) was added to the reaction buffer, and the mixture was incubated for another 30 min at 37°C to remove the phosphoryl group(s) of PYL2-GST or PYL4-GST. Primers used in this experiment are listed in Supplemental Table1.

### Cell-free degradation

To investigate the effects of GSO1 on PYLs, we determined PYLs-GST protein levels after incubation with total protein extracts from GSO1-OE-4, *gso1-1*, or WT seedlings grown on 1/2 MS medium with or without 1 μ M ABA. Total proteins were extracted from 0.4 g samples of 7-day-old seedlings of all genotypes using 600 ml of extraction buffer (5 mM dithiothreitol [DTT], 10 mM NaCl, 25 mM Tris–HCl [pH 7.5], 10 mM ATP, 4 mM phenylmethylsulfonyl fluoride, and 10 mM MgCl_2_). Crude extracts were held on ice (4°C) for 30 min and centrifuged twice at 12 000 rpm for 10 min at 4°C. After centrifugation, the supernatants were collected. About 0.5 mg of purified PYLs-GST was incubated with total protein extracts for 0, 20, 40, or 60 min at 22°C. At each time point, 20 ml of each solution was transferred to a new centrifuge tube, combined with 5 ml of loading buffer, and boiled for 5 min to terminate the reaction. The level of PYL protein in each reaction was determined using anti-PYL2 or PYL4 antibodies. Spot densitometry was performed using ImageJ v1.36 (http://rsb.info.nih.gov/ij/). To investigate the PYLs degradation pathway, PYLs protein expression level of 7-day-old WT, GSO1-OE and *gso1-1* seedlings over 4 h of 100 μM ABA plus MG132 (100 μM) treatment or 100 μM ABA plus E-64d (100 μM) before sampling. The total proteins were extracted and detected with anti-PYL2 or PYL4 antibodies. ImageJ v1.36 was used to quantify the intensity of each protein band.

### Stress treatments

For the inhibitor combined with drought treatment, seedlings were pre-treated with fluridone (50 μM) for 6 h and then placed on the filter paper for indicated times at 22°C. For ABA (100 μM) treatment, 7-day-old seedlings were sprayed with the configured ABA solution. Untreated seedlings were used as controls. Samples were collected after treatments at 0, 0.5, 3, 6 and 12 h for analysis.

The seedlings were collected and frozen in liquid nitrogen as soon as possible. Samples were collected in liquid nitrogen and stored at −80°C.

### Subcellular localization

For the observation of subcellular localization in the *Arabidopsis* roots, the 35Sp:GSO1-GFP construct was transformed into the *Agrobacterium* strain GV3101. The transformation method is similar as the BiFC mentioned above. 7-d-old 35Sp:GSO1-GFP seedlings were pre-treated with CHX (50 μM) for 1 h and then treated with CHX plus BFA (50 μM) or CHX plus BFA (50 μM) and 100 μM ABA for 0.5 h. Subsequently, the fluorescence signals in root tip cells were observed using Zeiss confocal microscopy (LSM880).

### LC-MS/MS for identification Phosphorylation of site of PYL2

For mass spectrometry analysis, protein samples were separated using SDS-PAGE electrophoresis and detected by coomassie brilliant blue staining. The SDS-PAGE gel at 47 kD was collected into a clean centrifuge tube. Cut the target bands into 1 mm³ gel particles and load them into 1.5 mL EP tubes. De-stain with a 50% ACN-50% 50 mmol/L NH ₄ HCO ₃ solution, incubate for 10-30 minutes, then pipette off and discard. Repeat this process until the gel particles are colorless. Add 1 mL of 100% ACN, let it stand for 30 minutes until the gel particles turn white and condense into clusters. Discard the ACN, then allow it to dry at room temperature. Add 10 mmol/L DTT at 100 μL per tube and reduce at 56°C in a water bath for 1 hour. Aspirate and discard. Add 55 mmol/L IAM at 100 μL per tube, react at room temperature in the dark for 1 hour. Aspirate and discard. Wash once with decolorization solution (ACN-50% 50 mmol/L NH_4_HCO_3_), aspirate and discard. Add 1000 μL of 100% ACN, incubate for 30 minutes until the gel particles turn white and condense into a clustered state. Discard the ACN and dry at room temperature. Take an enzyme with a concentration of 0.5 μg/μL (diluted with 100 mmol/L Tris HCl (pH 8.0)), add the enzyme solution at a rate of 7-10 μL per tube, incubate in a refrigerator at 4°C for 40 minutes, and then add 5-10 μL of 100 mmol/L Tris HCl (pH 8.0) solution to each tube. Seal and place in a 37°C water bath for enzyme digestion for 16 hours. Add 100 μL/tube of extraction solution (5% TFA-50% ACN-45% water), water bath at 37°C for 1 hour, sonicate for 5 minutes, centrifuge for 5 minutes, transfer the extraction solution to another new EP tube, repeat extraction once, combine the extraction solutions, and vacuum centrifuge dry. After enzymatic digestion, the peptide segments were desalinated using a self-filled desalination column and the solvent was evaporated in a 45°C vacuum centrifuge concentrator. The original mass spectrometry files were retrieved from the target protein database using Byonic, with the following search parameters: Fixed modifications: Carbamide (C), Variable modifications: Oxidation (M), Acetyl (N-term), Phosphorus (+79.966) (S and T), Enzymes: Chymotrypsin, Trypsin, Maximum Missed Cleavages: 3, First level mass spectrometry error (Peptide Mass Tolerance): 20 ppm, Fragment Mass Tolerance: 0.02Da.

### Statistics and reproducibility

Statistical analyses were performed using GraphPad Prism v.8 or Microsoft Excel 365 (v.16.0). Two-tailed paired t tests were performed when comparing two categories. Significance indicated as *P-value < 0.05, **P-value < 0.01, ***P-value <0.001 and “ns” means no significant difference. Apart from these, all remaining experiments were independently repeated at least three times, with consistent results.

## Data availability

All data supporting the findings of this study are available in the article, the extended data figures or the Supplementary information. Sequence data for the genes described in this article can be found in the *Arabidopsis* Information Resource (https://www.arabidopsis.org) under the following accession numbers: *GSO1* (AT4G20140), *PYR1* (AT4G17870), *PYL1* (AT5G46790), *PYL2* (AT2G26040), *PYL4* (AT2G38310), *PYL5* (AT5G05440), *PYL6* (AT2G40330), *PYL8* (AT5G53160), *PYL10* (AT4G27920), *PYL13* (AT4G18620), *IOS1* (AT1G51800), *CRLK2* (AT5G15730), *BRI1* (AT4G39400), *BAK1* (AT4G33430), *SARK1* (AT4G30520), *GHR1* (AT4G20940), *CIF1* (AT2G16385), *CIF2* (AT4G34600), *FREE1* (AT1G20110), *VPS23A* (AT3G12400).

## Acknowledgements

This work was supported by the Natural Science Foundation of China (32472064 and 32241039) and the Taishan Program of Shandong Province (grants tsqn202408127). We thank Dr. Chunming Liu and Xiufen Song (Key Laboratory of Plant Molecular Physiology, Institute of Botany, Chinese Academy of Sciences, Beijing, China) for providing seeds.

## Author contributions

C.-A.W., S.-Z. Z., and S.-K.G. conceived and designed the study. S.-K.G., Y.Z., L.Z., Q.-T.L., and J,-Y. L performed the experiments; C.-A.W. and S.-K.G. analyzed the data, made the figures, and wrote the manuscript; G.Y., S.Z., J.H., K.Y. and C.-C. Z provided experimental and writing suggestions. All authors approved the final manuscript.

## Supplementary data

**Supplemental Figure 1.**
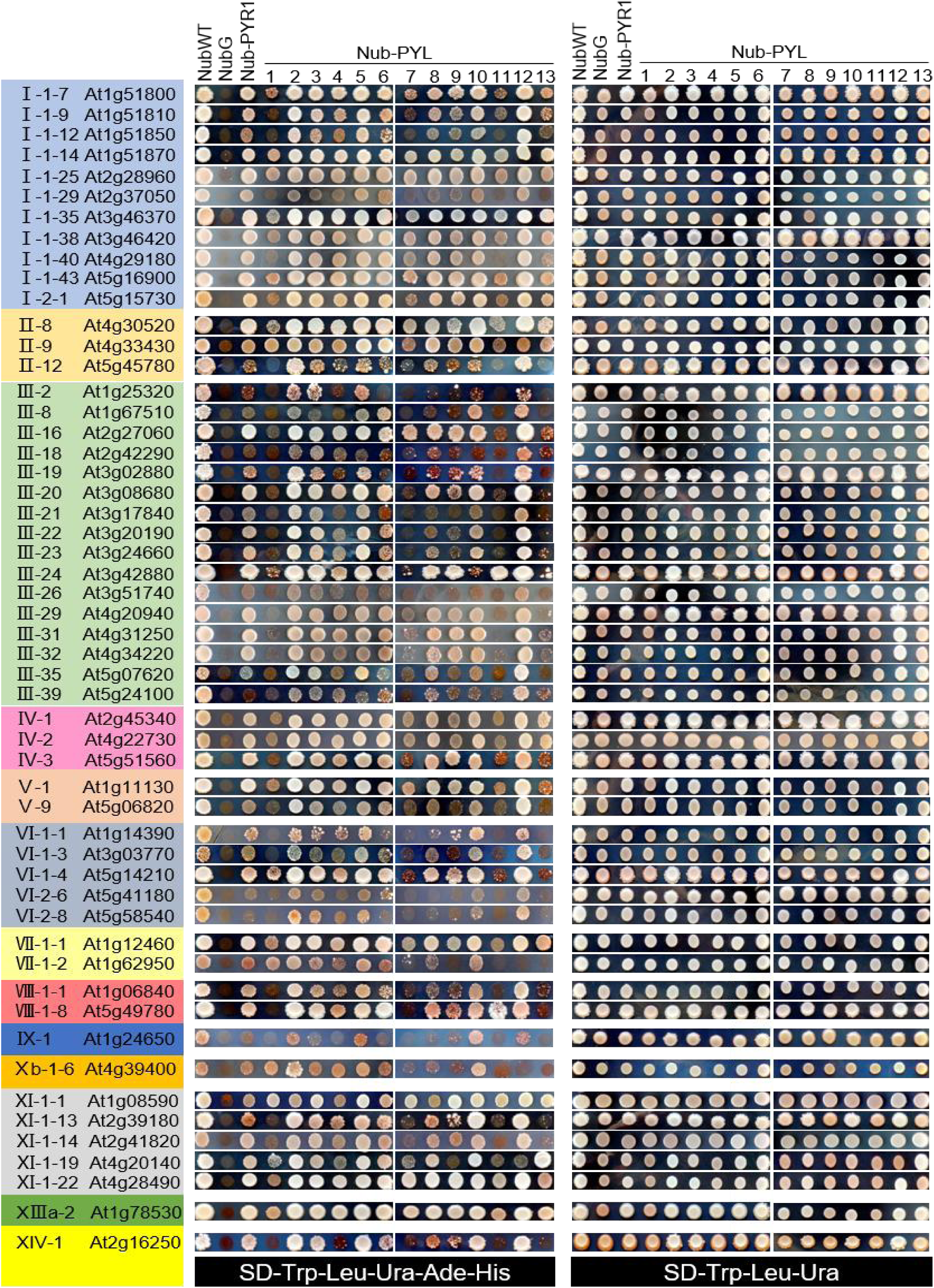
Membrane yeast two-hybrid (mbY2H) screening of interactions between 53 LRR-RLKs and 14 PYR/PYL receptors.

**Supplemental Figure 2.**
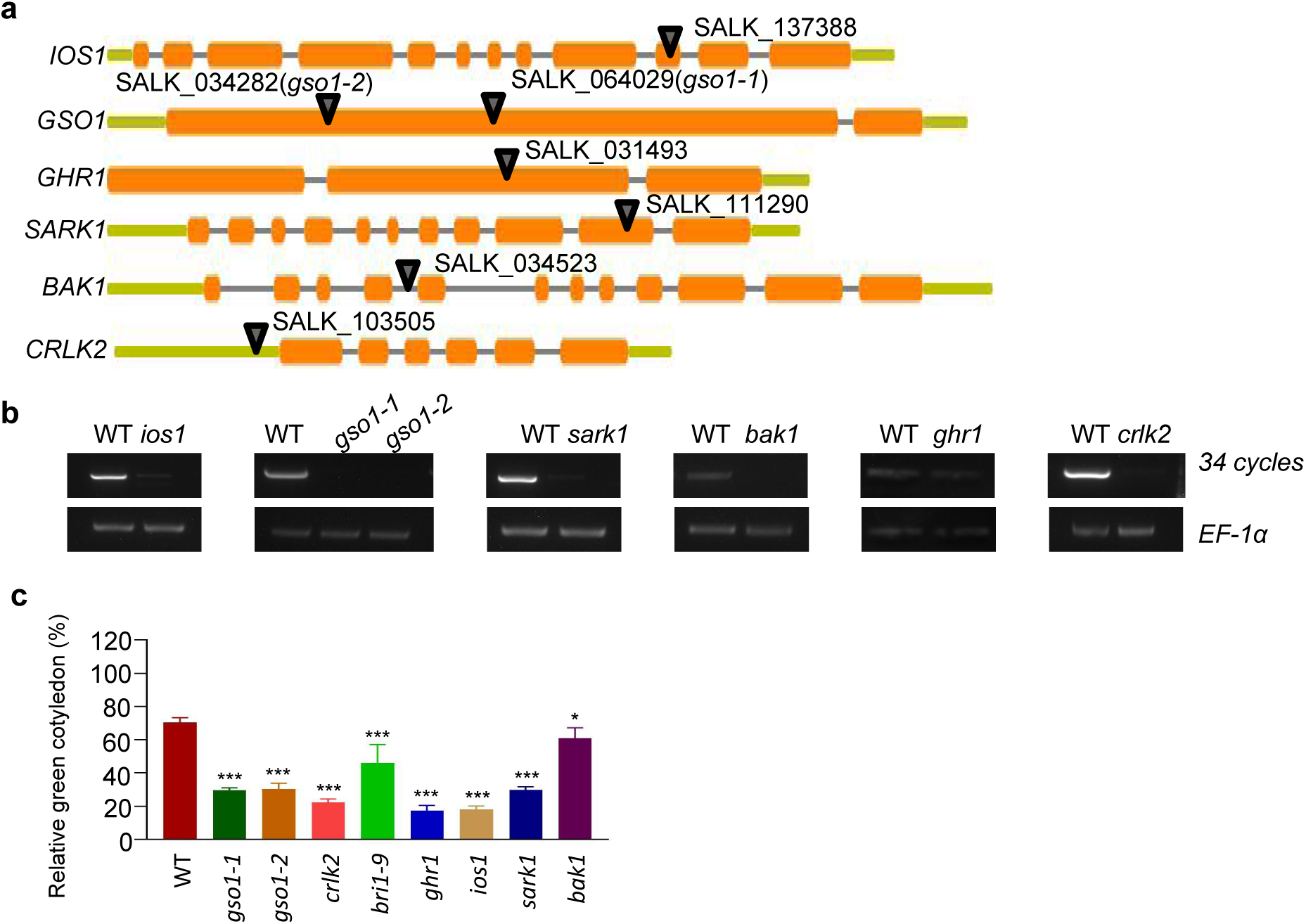
Identification and ABA responsive phenotypic analysis of seven *lrr-rlk* mutant lines. **a,** Schematic of T-DNA insertion sites in the seven *lrr-rlk* mutants. **b,** Transcription levels of the corresponding *LRR-RLK* genes in WT and mutant seedlings, detected by RT-PCR. *EF-1α* serves as the internal control. **c,** Relative green cotyledon rates of *Arabidopsis* seedlings in **Fig. 1b**. Data are mean ± SE of three independent experiments, each containing four technical replicates (n = 45 seedlings per replicate). Statistical significance was accessed by student’s *t*-test (*P < 0.05, **P < 0.01, ***P < 0.001).

**Supplemental Figure 3.**
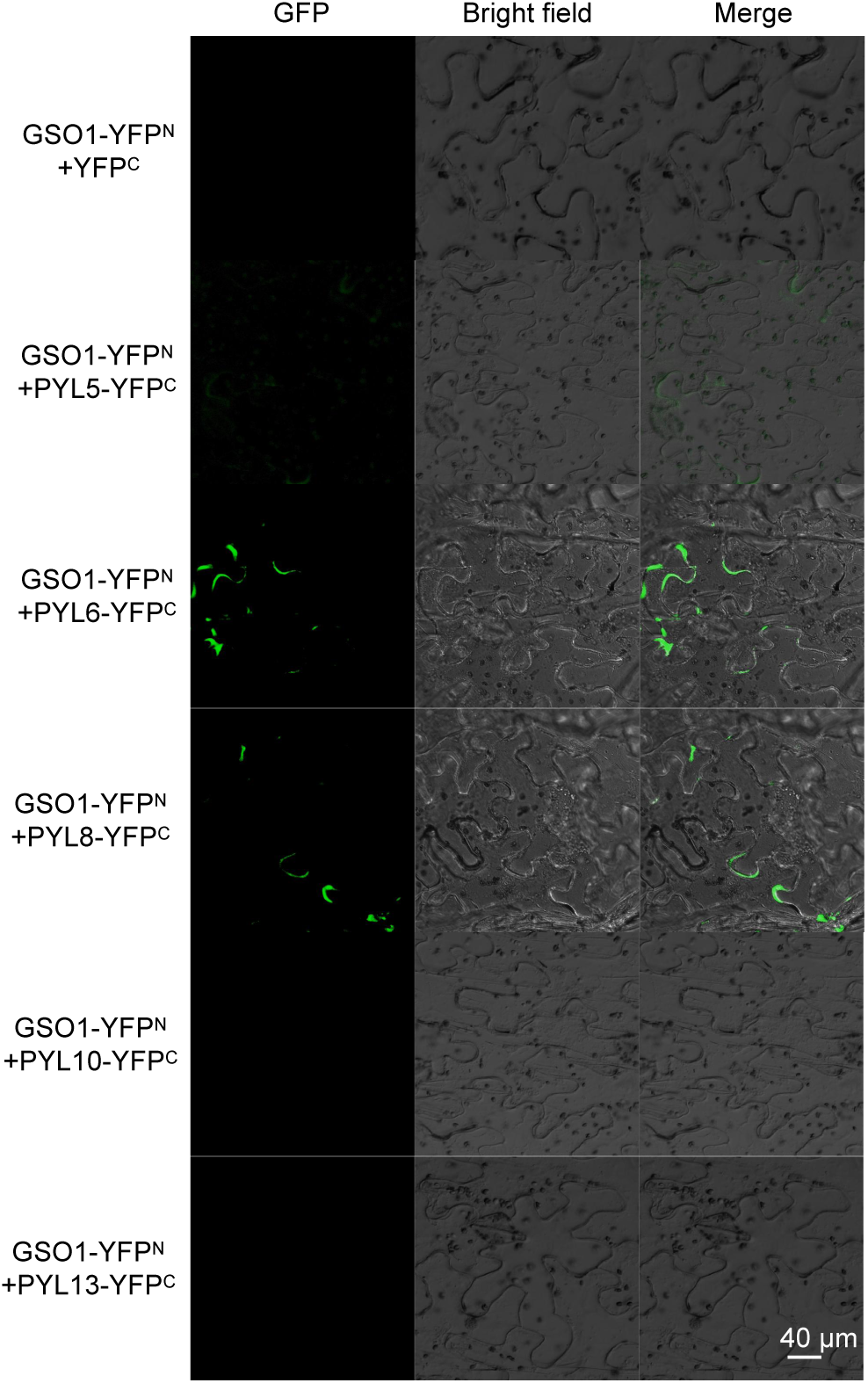
BiFC assays showing interactions between GSO1 and PYL5, PYL6, PYL8, PYL10, or PYL13 in *Nicotiana benthamiana*. Scale bar: 40 μm.

**Supplemental Figure 4.**
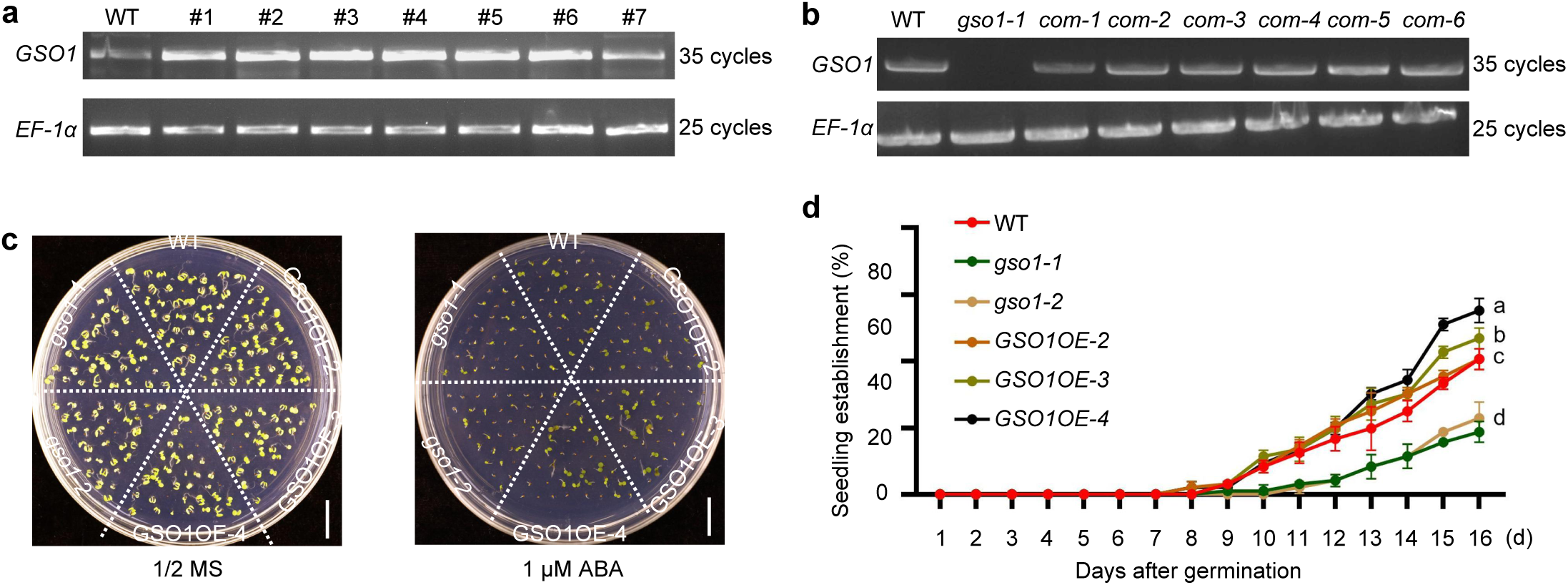
Identification and ABA-responsive phenotypic analysis of GSO1 overexpression and complementation lines. **a,** *GSO1* transcript levels in WT and *GSO1-* overexpressing lines, determined by RT–PCR. *EF-1a* was used as the internal control. **b,** *GSO1* transcript levels in WT, *gso1-1* and pGSO1::GSO1-GFP/*gso1-1* complementation lines, determined by RT–PCR. *EF-1a* was used as the internal control. **c,** Phenotypes of WT, *gso1-1*, *gso1-2* ,and GSO1-OE seedlings grown on 1/2 MS with or without 1 µM ABA for 10 d. Scale Bar: 1 cm. **d,** Seedling establishment rates of WT, *gso1-1*, *gso1-2* ,and GSO1-OE lines grown for 16 days on 1/2 MS medium with or without 1 μM ABA. Data are mean ± SEM (n = 3). Different lowercase letters indicate statistically significant differences (P < 0.05; one-way ANOVA).

**Supplemental Figure 5.**
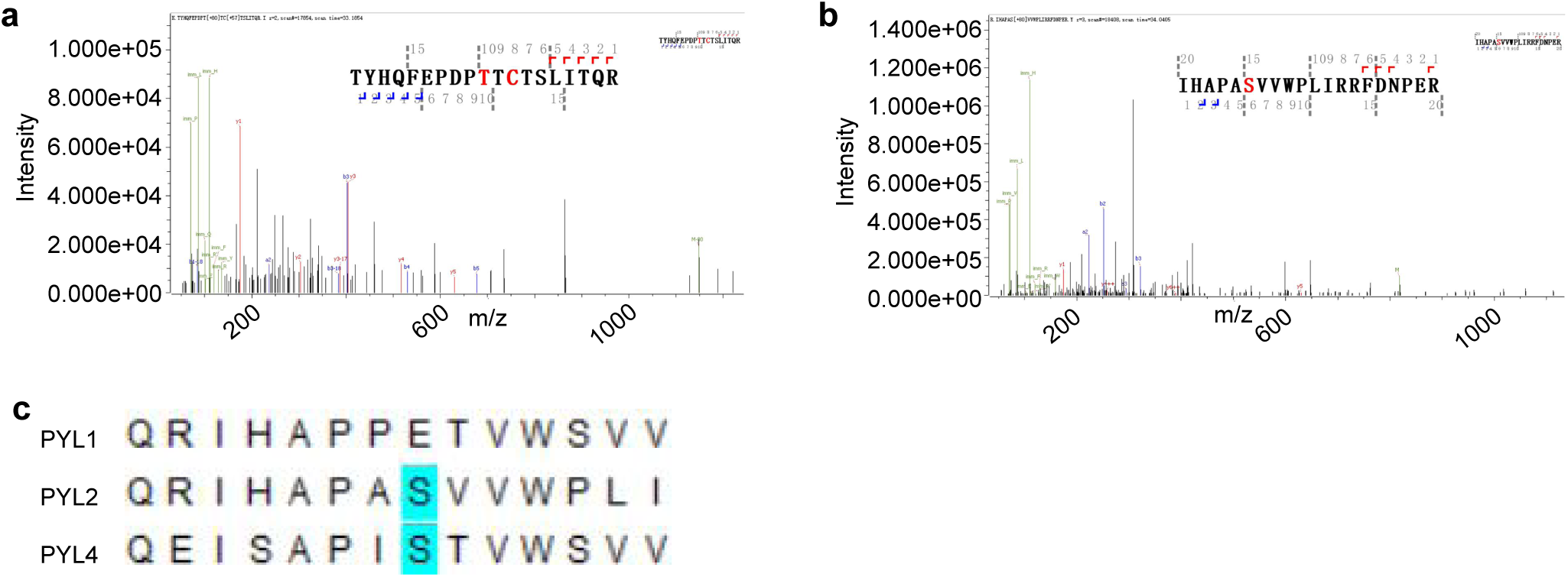
Identification of phosphorylation sites in PYL2 and PYL4. **a,b,** Phosphosites at Ser33 and Ser48 of PYL2 detected by liquid chromatography-mass spectrometry (LC-MS). Enriched phosphosites were analyzed by LC-MS. **c,** Multiple amino acid sequence alignment of PYL1, PYL2 , and PYL4. Conserved residues corresponding to PYL2 Ser33 and Ser48 are highlighted in blue boxes.

**Supplemental Figure 6.**
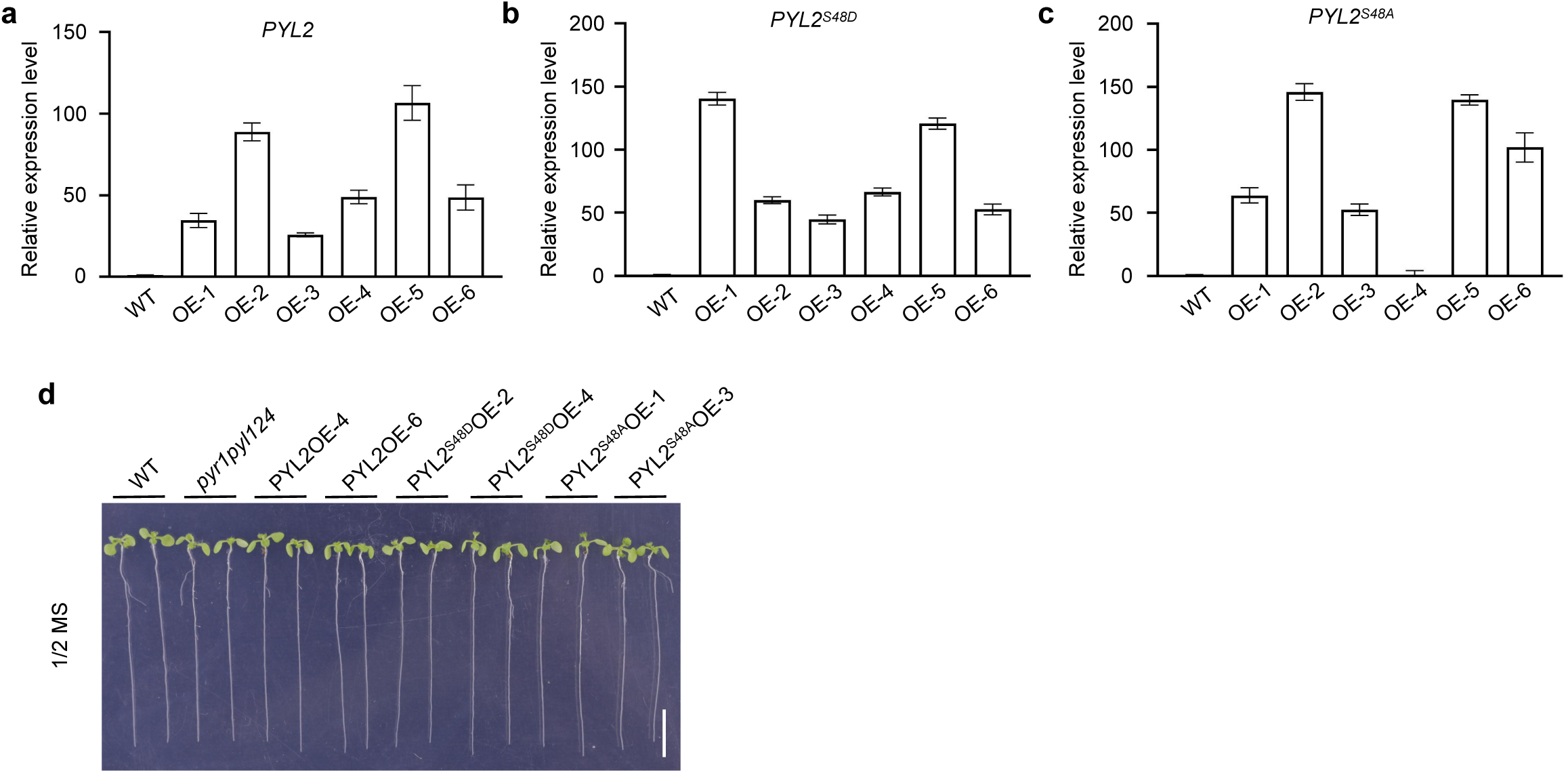
Characterization of the PYL2^WT^, PYL2^S48D, and^ PYL2^S48A^ overexpression lines. **a-c,** Transcript levels of *PYL2*, *PYL2^S48A^* ,and *PYL2^S48D^* in 7-day-old transgenic seedlings, determined by RT-qPCR. Expression levels were normalized to *18S* rRNA. Data are mean ± SEM (n = 3 biological replicates). Different lowercase letters indicate statistically significant differences (P < 0.05, one-way ANOVA). **d,** Phenotypes of 10-day-old WT, *pyr1pyl124*, PYL2OE, PYL2^S48A^OE, and PYL2^S48D^OE seedlings grown on 1/2 MS medium without ABA. Scale bar: 1 cm.

**Supplemental Figure 7.**
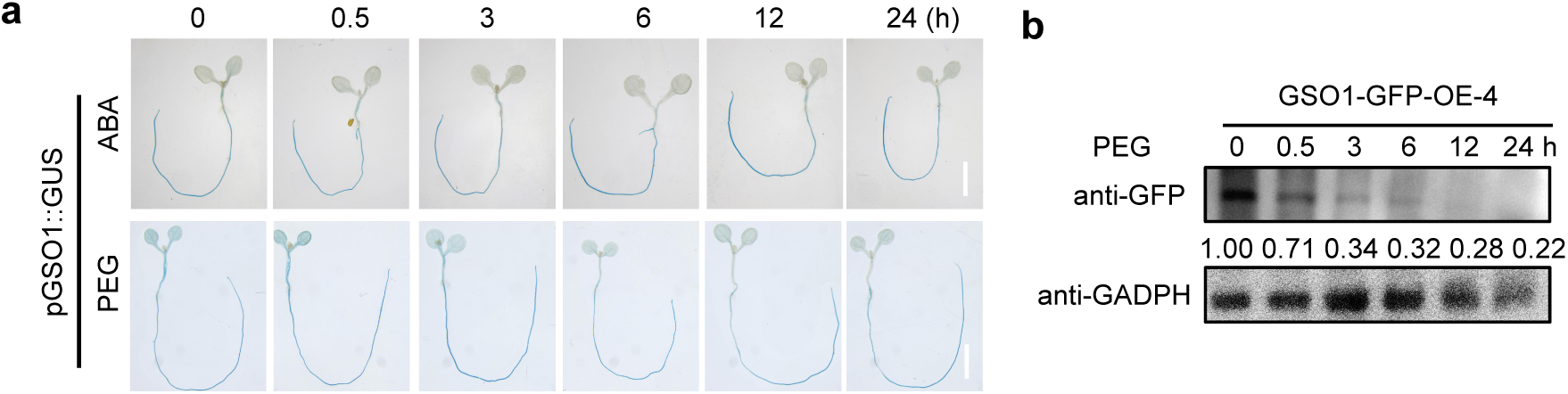
Effect of ABA and PEG on *GSO1* transcription and protein levels. **a,** GUS staining of *pGSO1::GUS* seedlings treated with 50 μM ABA or 30% PEG6000 for the indicated durations (0 - 24 h). Scale bar: 1cm. **b,** Stability of GSO1 protein under PEG treatment. Ten-day-old 35S:GSO1-GFP-OE-4 seedlings grown on 1/2 MS medium were treated with 30% PEG6000. GSO1-GFP was detected with anti-GFP antibody immunoblotting. GADPH serves as the loading control.

**Supplemental Figure 8.**
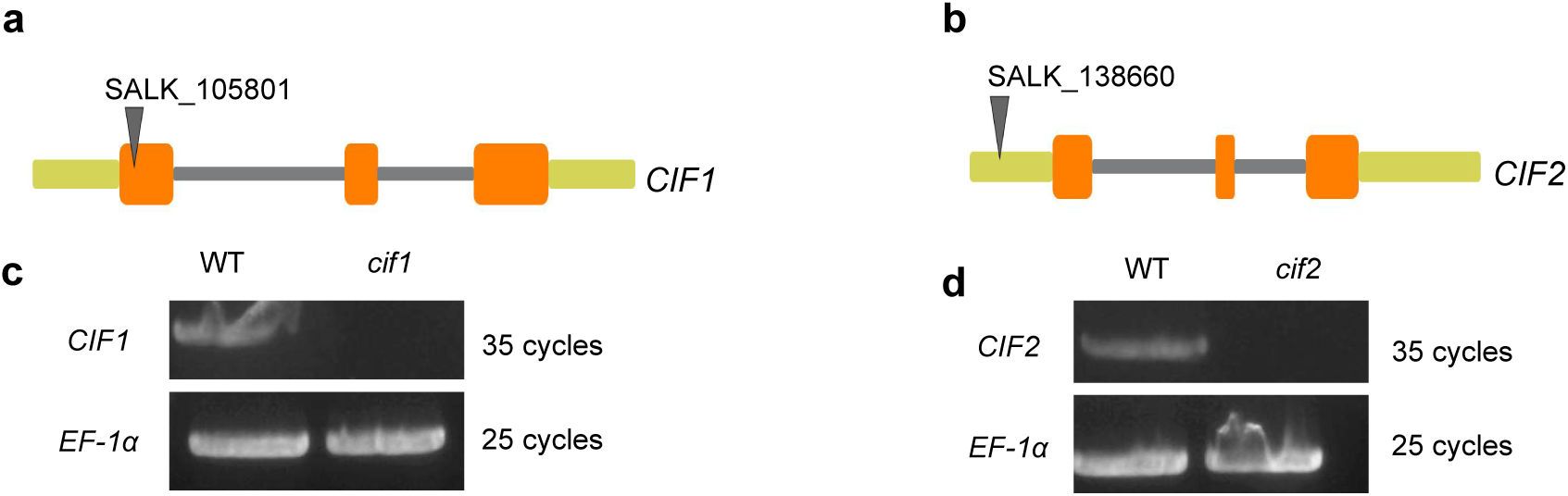
Characterization of *cif1* and *cif2* mutant lines. **a,b,** Schematic of T-DNA insertion sites in the *cif1* and *cif2* mutants. **c,d,** *CIF1* and *CIF2* transcript levels in WT, *cif1*, and *cif2* seedlings, analyzed by RT-PCR.

## References

1. Kuromori, T., Fujita, M., Takahashi, F., Yamaguchi-Shinozaki, K. & Shinozaki, K. Inter-tissue and inter-organ signaling in drought stress response and phenotyping of drought tolerance. Plant J. 109, 342–358 (2022).

2. Raghavendra, A.S., Gonugunta, V.K., Christmann, A. & Grill, E. ABA perception and signalling. Trends Plant Sci. 15, 395–401 (2010).

3. Lee, S.C. & Luan, S. ABA signal transduction at the crossroad of biotic and abiotic stress responses. Plant Cell Environ. 35, 53–60 (2012).

4. Cutler, S.R., Rodriguez, P.L., Finkelstein, R.R. & Abrams, S.R. Abscisic Acid: Emergence of a Core Signaling Network. Annu. Rev. Plant Biol. 61, 651–679 (2010).

5. Fujii, H. et al. In vitro reconstitution of an abscisic acid signalling pathway. Nature 462, 660–U138 (2009).

6. Gonzalez-Guzman, M. et al. Arabidopsis PYR/PYL/RCAR Receptors Play a Major Role in Quantitative Regulation of Stomatal Aperture and Transcriptional Response to Abscisic Acid. Plant Cell 24, 2483–2496 (2012).

7. Park, S.Y. et al. Abscisic Acid Inhibits Type 2C Protein Phosphatases via the PYR/PYL Family of START Proteins. Science 324, 1068–1071 (2009).

8. Ma, Y. et al. Regulators of PP2C Phosphatase Activity Function as Abscisic Acid Sensors. Science 324, 1064–1068 (2009).

9. Chong, L., Xu, R., Ku, L. & Zhu, Y. Beyond stress response: OST1 opening doors for plants to grow. Stress Biology 2(2022).

10. Zhao, Y. et al. The ABA Receptor PYL8 Promotes Lateral Root Growth by Enhancing MYB77-Dependent Transcription of Auxin-Responsive Genes. Science Signaling 7(2014).

11. Lu, C.T. et al. dbPTM 3.0: an informative resource for investigating substrate site specificity and functional association of protein post-translational modifications. Nucleic Acids Res. 41, D295–D305 (2013).

12. Yu, Z.P. et al. CEPR2 phosphorylates and accelerates the degradation of PYR/PYLs in Arabidopsis. J. Exp. Bot. 70, 5457–5469 (2019).

13. Zhang, L. et al. SiCEP3, a C-terminally encoded peptide from Setaria italica, promotes ABA import and signaling. J. Exp. Bot. 72, 6260–6273 (2021).

14. Diévart, A. & Clark, S.E. Using mutant alleles to determine the structure and function of leucine-rich repeat receptor-like kinases. Curr Opin Plant Biol 6, 507–16 (2003).

15. Ogawa, M., Shinohara, H., Sakagami, Y. & Matsubayashi, Y. Arabidopsis CLV3 peptide directly binds CLV1 ectodomain. Science 319, 294 (2008).

16. Hu, C. et al. A group of receptor kinases are essential for CLAVATA signalling to maintain stem cell homeostasis. Nat. Plants 4, 205–211 (2018).

17. Nam, K.H. & Li, J. BRI1/BAK1, a receptor kinase pair mediating brassinosteroid signaling. Cell 110, 203–12 (2002).

18. Song, W.Y. et al. A receptor kinase-like protein encoded by the rice disease resistance gene, Xa21. Science 270, 1804–1806 (1995).

19. Giordano, L., Schimmerling, M., Panabières, F., Allasia, V. & Keller, H. The exodomain of the impaired oomycete susceptibility 1 receptor mediates both endoplasmic reticulum stress responses and abscisic acid signalling during downy mildew infection of Arabidopsis. Mol. Plant Pathol. 23, 1783–1791 (2022).

20. Hok, S. et al. The Receptor Kinase IMPAIRED OOMYCETE SUSCEPTIBILITY1 Attenuates Abscisic Acid Responses in Arabidopsis. Plant Physiol. 166, 1506–1518 (2014).

21. Sierla, M. et al. The Receptor-like Pseudokinase GHR1 Is Required for Stomatal Closure. Plant Cell 30, 2813–2837 (2018).

22. Yang, Y. et al. The phospholipid flippase ALA3 regulates pollen tube growth and guidance in Arabidopsis. Plant Cell 34, 3718–3736 (2022).

23. Liu, M.L. et al. AtLURE1/PRK6-mediated signaling promotes conspecific micropylar pollen tube guidance. Plant Physiol. 186, 865–873 (2021).

24. Xiang, X.J. et al. Arabidopsis class A S-acyl transferases modify the pollen receptors LIP1 and PRK1 to regulate pollen tube guidance. Plant Cell 36, 3419–3434 (2024).

25. Tsuwamoto, R., Fukuoka, H. & Takahata, Y. GASSHO1 and GASSHO2 encoding a putative leucine - rich repeat transmembrane - type receptor kinase are essential for the normal development of the epidermal surface in Arabidopsis embryos. Plant J. 54, 30–42 (2007).

26. Nakayama, T. et al. A peptide hormone required for Casparian strip diffusion barrier formation in Arabidopsis roots. Science 355, 284–286 (2017).

27. Creff, A. et al. A stress-response-related inter-compartmental signalling pathway regulates embryonic cuticle integrity in Arabidopsis. PLoS Genet. 15(2019).

28. Zhang, H. et al. SERKs regulate embryonic cuticle integrity through the TWS1-GSO1/2 signaling pathway in Arabidopsis. N. Phytol. 233, 313–328 (2022).

29. Okuda, S. et al. Molecular mechanism for the recognition of sequence-divergent CIF peptides by the plant receptor kinases GSO1/SGN3 and GSO2. Proc. Natl Acad. Sci. USA 117, 2693–2703 (2020).

30. Racolta, A., Bryan, A.C. & Tax, F.E. The Receptor-Like Kinases GSO1 and GSO2 Together Regulate Root Growth in Arabidopsis Through Control of Cell Division and Cell Fate Specification. Dev. Dyn. 243, 257–278 (2014).

31. Chen, X.X. et al. Protein kinases in plant responses to drought, salt, and cold stress. J. Integr. Plant Biol. 63, 53–78 (2021).

32. Ronkina, N. & Gaestel, M. MAPK-Activated Protein Kinases: Servant or Partner? Annu. Rev. Biochem. 91, 505–540 (2022).

33. Isono, E. & Kalinowska, K. ESCRT-dependent degradation of ubiquitylated plasma membrane proteins in plants. Curr Opin Plant Biol 40, 49–55 (2017).

34. Gao, C.J., Zhuang, X.H., Shen, J.B. & Jiang, L.W. Plant ESCRT Complexes: Moving Beyond Endosomal Sorting. Trends Plant Sci. 22, 986–998 (2017).

35. Luschnig, C. & Vert, G. The dynamics of plant plasma membrane proteins: PINs and beyond. Development 141, 2924–2938 (2014).

36. Shiu, S.H. & Bleecker, A.B. Receptor-like kinases from Arabidopsis form a monophyletic gene family related to animal receptor kinases. Proc. Natl Acad. Sci. USA 98, 10763–10768 (2001).

37. Razi, K. & Muneer, S. Drought stress-induced physiological mechanisms, signaling pathways and molecular response of chloroplasts in common vegetable crops. Crit. Rev. Biotechnol. 41, 669–691 (2021).

38. Gupta, A., Rico-Medina, A. & Caño-Delgado, A.I. The physiology of plant responses to drought. Science 368, 266–269 (2020).

39. Gong, Z.Z. et al. Plant abiotic stress response and nutrient use efficiency. SCI CHINA LIFE SCI 63, 635–674 (2020).

40. Chen, H.H., Qu, L., Xu, Z.H., Zhu, J.K. & Xue, H.W. EL1-like Casein Kinases Suppress ABA Signaling and Responses by Phosphorylating and Destabilizing the ABA Receptors PYR/PYLs in Arabidopsis. Mol. Plant 11, 706–719 (2018).

41. You, Z. et al. The CBL1/9-CIPK1 calcium sensor negatively regulates drought stress by phosphorylating the PYLs ABA receptor. Nat. Commun. 14(2023).

42. Liu, S.S. et al. PLATZ2 negatively regulates salt tolerance in Arabidopsis seedlings by directly suppressing the expression of the CBL4/SOS3 and CBL10/SCaBP8 genes. J. Exp. Bot. 71, 5589–5602 (2020).

43. Liu, L. et al. Function identification of MdTIR1 in apple root growth benefited from the predicted MdPPI network. J. Integr. Plant Biol. 63, 723–736 (2021).

44. Szklarczyk, D. et al. The STRING database in 2023: protein–protein association networks and functional enrichment analyses for any sequenced genome of interest. Nucleic Acids Res. 51, D638–D646 (2023).

45. Smoot, M.E., Ono, K., Ruscheinski, J., Wang, P.-L. & Ideker, T. Cytoscape 2.8: new features for data integration and network visualization. Bioinformatics 27, 431–432 (2011).

46. Chen, C.T. et al. Arabidopsis SAG protein containing the MDN1 domain participates in seed germination and seedling development by negatively regulating ABI3 and ABI5. J. Exp. Bot. 65, 35–45 (2014).

47. Chen, H. et al. Firefly Luciferase Complementation Imaging Assay for Protein-Protein Interactions in Plants. Plant Physiol. 146, 323–324 (2008).

48. Waadt, R. & Kudla, J. In Planta Visualization of Protein Interactions Using Bimolecular Fluorescence Complementation (BiFC). Cold Spring Harbor Protocols 2008(2008).

